# Remodeling of human colon plasma cell repertoire in ulcerative colitis

**DOI:** 10.1101/2022.02.14.480403

**Authors:** Johannes F. Scheid, Basak Eraslan, Andrew Hudak, Eric Brown, Dallis Sergio, Toni Delorey, Devan Phillips, Ariel Lefkovith, Alison T. Jess, Lennard W. Duck, Charles O. Elson, Hera Vlamakis, Jacques Deguine, Ashwin Ananthakrishnan, Daniel B. Graham, Aviv Regev, Ramnik J. Xavier

**Author notes:** These authors contributed equally (Johannes F. Scheid, Basak Eraslan).

## Abstract

Plasma cells (PCs) constitute a significant fraction of cells in colonic mucosa and contribute to inflammatory lymphocytic infiltrates in ulcerative colitis (UC). While gut PCs secrete 3-5 g of immunoglobulins daily, including IgA antibodies that target colitogenic bacteria, their role in UC is not known. Here, we combined B cell sorting with single-cell VDJ- and RNA-seq and monoclonal antibody (mAb) testing to characterize the colonic PC repertoire in healthy individuals and patients with UC. We show that a large fraction of B cell clones is shared between different colon regions and that inflammation in UC disrupts this landscape, causing clonal expansion and isotype skewing from IgA1 and IgA2 to IgG1. mAbs produced from expanded PC clones show low polyreactivity and autoreactivity and target specific bacterial strains. Expression profiles of individual PCs from inflamed and non-inflamed colon regions indicate that inflammation is associated with up-regulation of the unfolded protein response (UPR) and antigen presentation genes. Together, our results characterize the microbiome-specific PC response in the colon, its disruption in UC and how PCs might contribute to inflammation in UC.

## INTRODUCTION

Despite advances in ulcerative colitis (UC) treatment many patients with poorly controlled disease still have to undergo colectomy^1^ and a better understanding of the disease is needed in order to identify new treatment targets. In particular, the role of microbiome-directed adaptive immunity in UC remains elusive^2^. Plasma cells (PCs) constitute a major fraction of lymphocytes in UC-associated inflammatory infiltrates and their expansion is linked to the risk of disease recurrence^3^. In addition, an Fc receptor variant (FcgR2) with decreased affinity to IgG1 is protective against UC in GWAS studies suggesting a role for IgG mAbs in the pathogenesis of UC^3, 4^. PCs in the human gut secrete grams of mAbs daily into the intestinal lumen^5^ and some of these are thought to target colitogenic bacteria^6^. However, the composition of the PC repertoire throughout the human colon in health and UC remains elusive.

Here, we used single cell VDJ- and RNA-seq profiling (sc(VDJ+RNA)-Seq) of healthy control subjects (HCs), UC patients in remission and those with active inflammation, to conduct an in-depth analysis of the mAb repertoire and PC expression states throughout the human colon. We show that inflammation is associated with a shift from IgA to IgG, disruption of PC clonal architecture, upregulation of the unfolded protein response (UPR) and antigen presentation genes in PCs, and that IgG1 mAbs from inflamed colon PCs specifically target pathogenic bacterial strains.

## RESULTS

### Single-cell profiling and VDJ characterization of human colon PCs in UC

To understand the effect of UC on the colonic PC repertoire, we profiled by sc(VDJ+RNA)-Seq 181,124 PCs from 49 colon biopsies or mucosal resection samples from 8 UC patients with at least one area of mucosal inflammation (12 inflamed and 9 less inflamed samples from the same subjects), 5 UC patients in endoscopic and histologic remission (8 non-inflamed samples) and 8 HCs (20 healthy samples) (Fig. 1a, **Methods**). The 21 subjects had equal representation of gender (10 women, 11 men) and ranged in age from 28 to 75 years; UC patients were diagnosed 1-20 years prior to enrollment and were on different therapeutic regimens (Extended Data Table 1). We digested colon samples to single cell suspensions, performed fluorescence-activated cell sorting (FACS) of CD27+CD38+ cells (5-20% of all cells, Fig. 1b) and profiled sorted cells by 5’ directed single-cell RNA-Seq (scRNA-seq) for both mRNA and paired VDJ profiling (sc(VDJ+RNA)-Seq), recovering matched single-cell VDJ and expression profiles, yielding 2,237-16,802 high quality cells per donor (Fig. 1c and Extended Data Fig. 1a, **Methods**).

**Figure 1:**
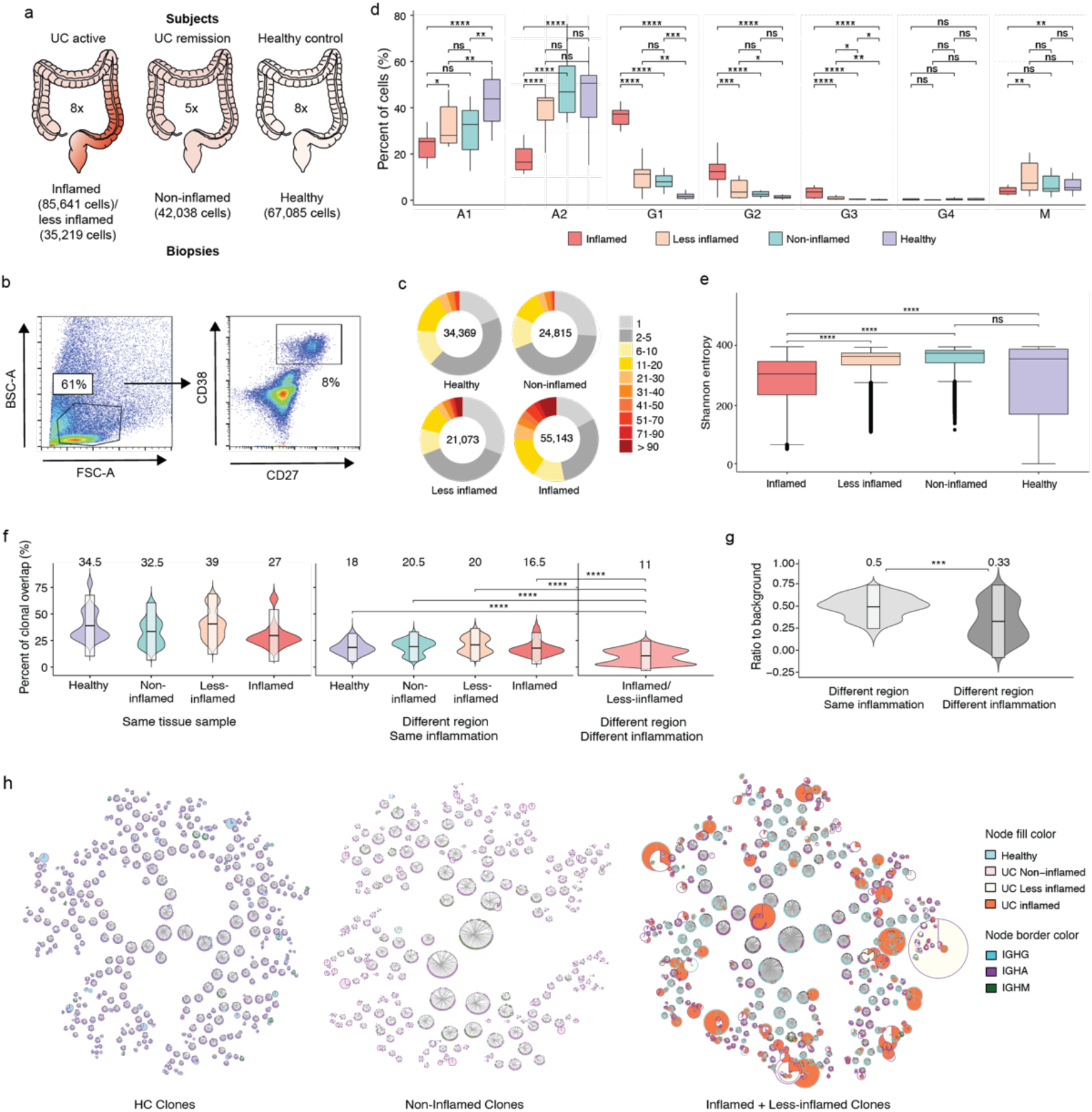
Patient selection, PC sorting, isotype analysis and clonal landscape. **a,** Three patient categories recruited in this study with the number of subjects indicated in the center, respectively. Left: UC patients with inflammation, center: UC patients in remission and right: HCs. The number of cells included in transcriptome analysis is indicated for each sample type. **b,** Representative FACS plots with the gating strategy for sorting of colon PCs. Cells isolated from digestion of colon biopsies and resection samples were gated on live cells based on their appearance in side scatter (BSC) and forward scatter (FSC). Of these, CD38-FITC and CD27-PE double-positive cells were selected for sorting and sequencing. **c,** Pie charts show the expansion of differently sized PC clones for all samples grouped based on their inflammation status. Numbers in the center of the pie charts stand for the total number of cells analyzed in that particular plot. **d,** Box-plots displaying the distribution of the percentage of immunoglobulin isotypes (y-axis) across samples grouped by their inflammation status (x-axis). Brackets indicate statistical significance using a one-sided non-parametric Wilcoxon test with *P ≤ 0.05, **P ≤ 0.01, ***P ≤ 0.001, ****P ≤ 0.0001 and n/s indicating no statistical significance. **e**, Box plots displaying the distribution of the Shannon entropy values of the samples stratified by their inflammation status as indicated. Shannon entropy is a measure of population diversity which is reversely related with the clonal expansion (**Methods**). Brackets indicate statistical significance using a one-sided t-test with \*\*\*\**P ≤* 0.0001 and n/s indicating no statistical significance. **f**, Violin plots displaying the distribution of the percentage of the shared clones between randomly sampled sets of PCs (**Methods**). Here the two random samples of a specific donor that are to be evaluated for clonal overlap can belong to i) same tissue sample (left), ii) different colon region with same inflammation status (center) and iii) different colon region with different inflammation status (right). The overlap between random samples that belong to different colon regions with the same inflammation status is significantly smaller (P value <= 0.0001, one-sided t-test) than the random samples that belong to the same colon regions for all healthy, non-inflamed, less inflamed and inflamed groups. The overlap reduces significantly (P value <= 0.0001, one-sided t-test) when the compared samples differ in their inflammation status. The boxes represent −2 standard deviation, mean, and + 2 standard deviation. The values above each violin plot represent the median values of the distribution. **g**, Distribution of the clonal overlap between samples of PCs that are collected from different colon regions corrected for the background overlap percentages of the parent regions (**Methods**). Bracket indicates statistical significance using a one-sided t-test with \*\*\**P ≤* 0.001. The boxes represent −2 standard deviation, mean, and + 2 standard deviation. The values above each violin plot represent the median values of the distribution. **h**, Schematic representation of the B cell clones in healthy, non-inflamed and inflamed samples, as indicated, show the clonal expansion and the isotype change in UC patients. Up to 50 of the largest clones from each donor are displayed. Each ring shows one distinct clone where the cells of the clone are depicted as individual nodes lined at the border of the clone ring and connected to each other by lines. The sizes of the nodes correspond to the number of the cells with the identical VDJ heavy chain sequence. The border color of the nodes stands for the Ig isotype and the fill colors of the nodes indicate the inflammation status of the samples these particular cells belong to.

### A shift from IgA1/A2 to IgG1 PCs in inflammation

Consistent with early immunohistochemistry analyses^7^ and prior single-cell atlases of the colon^8^, a significant skewing from IgA1 and IgA2 toward IgG1 occurred in inflamed samples, and to a smaller degree in less inflamed and non-inflamed samples, when compared to HCs (Fig. 1d, **Methods**, tested with both multivariate test accounting for compositional dependencies and one-sided non-parametric Wilcoxon rank-sum test, P-value < 10^-4^). IgG1 was the most frequent isotype in PCs from inflamed samples, but IgG2 and IgG3 were also significantly elevated when compared to PCs from HCs (Fig. 1d and Extended Data Fig. 1b, **Methods,** tested with both multivariate test accounting for compositional dependencies and one-sided non-parametric Wilcoxon rank-sum test, P-value < 10^-4^). Thus, PCs in UC are significantly skewed toward IgG1 when compared to HCs and this difference is most pronounced in inflamed tissue samples.

### Clonal expansion, disruption of clonal architecture, and shift from Igk to Igl light chain usage in PCs from inflamed colon

Most (56-86%) B cells from the 21 subjects belonged to clones of three or more cells based on their VDJ transcripts (Extended Data Fig. 1c). As an indication of significant clonal expansion Shannon entropy and Simpson index decreased in PCs from inflamed biopsies compared to all other disease states (HCs, uninflamed biopsies from remission, and less inflamed biopsies from subjects with active disease) (Fig. 1e, Extended Data Fig. 1c, **inverse Simpson index not shown**). Most of the 560 larger expanded clones (clone size > 9 cells) that were present in both inflamed and matched less inflamed colon areas from the same patient showed either IgG (166 of 560 clones) or IgA (375 of 560) dominance in inflamed areas or IgA (409 out of 560) dominance in the less inflamed colon areas, and significantly maintained their dominant isotype across regions with different inflammation status (Spearman’s ρ = 0.4, P-value < 2.2*10^-16^, Extended Data Fig. 1d, e, **Methods**).

Estimating clonal overlap between randomly selected sets of 100 cells from the same sample or different colon regions within a patient (**Methods**), the largest overlap was found among PCs within the same sample, followed by separate colon regions with the same inflammation state, where overlap was significantly higher than for regions with different inflammation states (Fig. 1f, g and Extended Data Fig. 2a, b, **Methods,** one tailed t-test, P-value <= 10^-3^). This held true when expanded clones were collapsed to one cell to correct for different levels of clonal expansion (Extended Data Fig. 2c, **Methods**). Identical clonal members (defined as a clonal member carrying the same heavy chain nucleotide sequence) were also significantly expanded in inflamed samples (Fig. 1h). Although PCs expressing Igk light chains were more prevalent than Igl light chains in all samples, there was a significant shift from Igk to Igl usage in UC samples (inflamed, less inflamed or non-inflamed) compared to HCs (Extended Data Fig. 3a, one-sided non-parametric Wilcoxon rank-sum test, P-value < 10^-4^). VH and VL gene repertoires were consistent across antibody isotypes and disease states (Extended Data Fig. 3b-d, **Methods**). Other antibody features such as CDRH3 length, charge and hydrophobicity, or inferred selection pressure (**Methods**) differed across antibody isotypes and disease states, some reaching statistical significance (Extended Data Fig. 3e-h, **Methods**). Taken together, inflammation in UC is associated with increased clonal PC expansion and clonal overlap between separate regions with shared inflammation status.

### Cell intrinsic expression changes in colon PCs during inflammation

The scRNA-seq profiles of the 181,124 sorted PCs partitioned into four main clusters (TCs), with one cluster (cluster 0) shared across all subjects (Fig. 2a and Extended Data Fig. 4a-c). Marker genes for this cluster include *NR4A1* and *EZR* which are both involved in the response to B cell receptor-antigen interactions^9, 10^(Extended Data Fig. 4b). Cluster 1, with marker genes *HSPA1A* and *HSPB1* that are upregulated in cellular stress^11^, and cluster 3, with marker genes *DERL3* and *SLC3A2* that are involved in the UPR^12, 13^, were specific to a small number of subjects (c, Extended Data Fig. 4c); cluster 2 was enriched in cells in cell cycle phases G2 and S (Fig. 2b). We observed an apparent separation between cells from inflamed vs. all other samples (Fig. 2c), even when restricting only to cells from subjects with matched inflamed and less inflamed samples (Extended Data Fig. 4d, e). Consistent with our PC isotype analysis above, IgG was enriched in clusters associated with inflammation (Fig. 2d).

**Figure 2:**
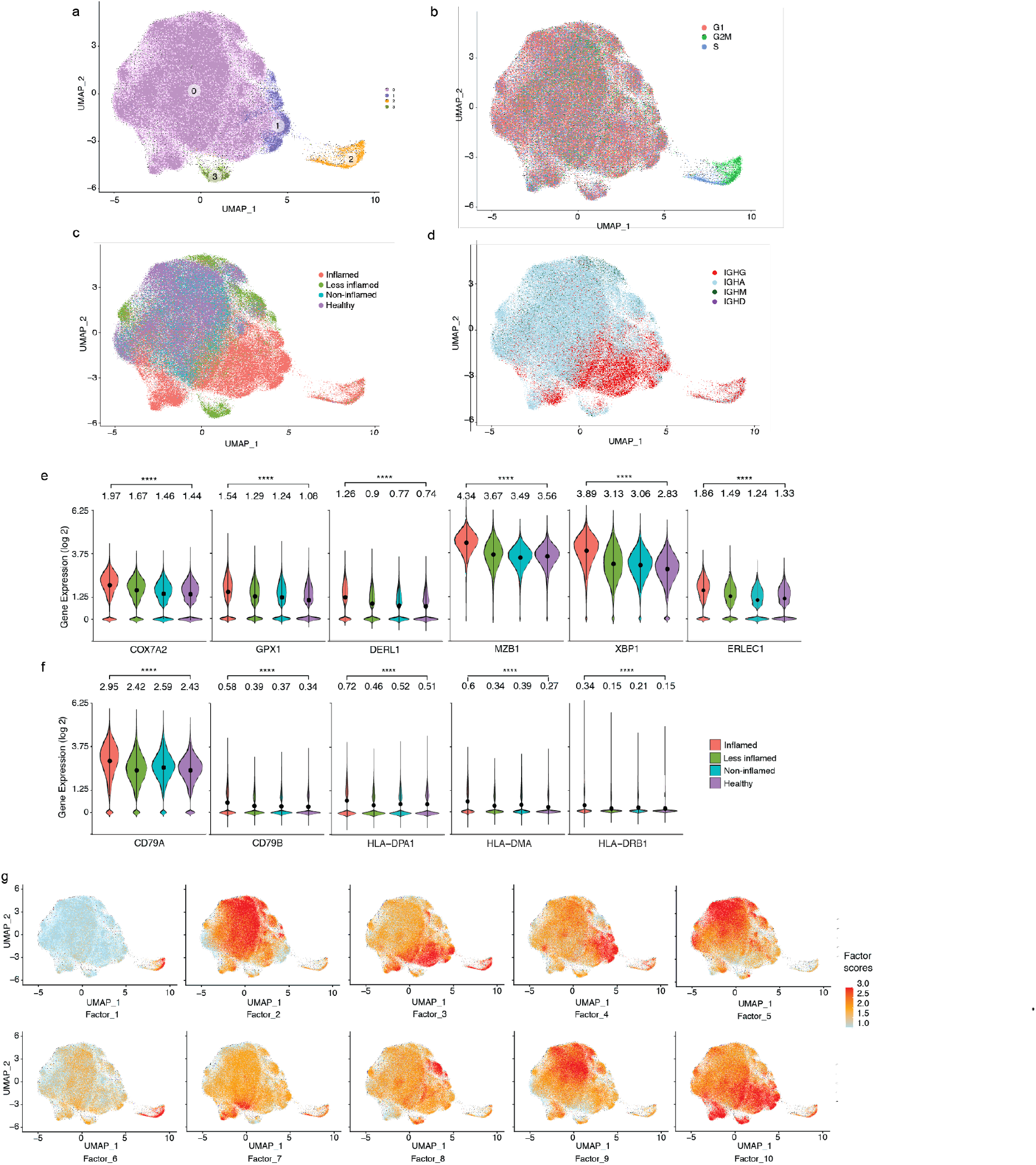
Transcriptional characteristics of PCs from inflamed colon areas. **a,** UMAP plot showing the cell embeddings based on the transcriptome. PCs are grouped into 4 clusters based on their transcriptomes. **b,** Same UMAP as in (**a**), but cells are colored based on their cell cycle phase G1 (red), G2M (green) or S (blue) as predicted by the cellular expression levels of the cell cycle genes^45, 46^. **c, d,** UMAP plots showing the cell embeddings colored by the (**c**) inflammation status of the tissue the cells were isolated from and (**d**) the antibody isotype of the cells. **e, f,** Selected example genes involved in oxidative stress and UPR-pathway (**e**) and antigen presentation (**f**) that are significantly differentially expressed (pseudobulk DE analysis, FDR < 0.05) between UC Inflamed and healthy samples. The violin plots display the distribution of the gene expression values per cell. Stars indicate statistical significance of the non-parametric Wilcoxon rank sum test. The values above each violin plot and dots in each violin plot indicate the mean value of the distribution. (**g**) Latent factors that generate the main sources of variation in the transcriptome. We identified 10 latent factors that generate the variation in the transcriptome of the investigated colon PCs (Extended Data Fig. 6). The 10 UMAP plots display the cells colored by the factor scores (**Methods**) for each of the 10 factors.

Genes differentially upregulated in PCs from inflamed compared to healthy colon areas across all cells (Extended Data Fig. 5a, **Methods,** pseudobulk DE analysis, FDR < 0.05) included: *COX7A2,* upregulated in antibody-secreting PCs undergoing oxidative metabolism^14^; *GPX1,* encoding an intracellular antioxidant enzyme upregulated in response to oxidative stress^15–17^; and endoplasmic reticulum (ER) stress-related genes^18–21^ *DERL1*, *MZB1*, *XBP1*, *ERLEC1* and *TMBIM6,* consistent with increased ER expansion to meet the demands of immunoglobulin secretion in PCs from inflamed areas (Fig. 2e, pseudobulk DE analysis, FDR < 0.05, Extended Data Fig. 5a, **Supplementary Table 1)**. Several genes involved in antigen uptake and presentation (*CD79A*, *CD79B* and several MHCII genes) were upregulated in PCs from inflamed areas (Fig. 2f, pseudobulk DE analysis, FDR < 0.05, Extended Data Fig. 5b,c), possibly mirroring the role of PCs in celiac disease as antigen presenting cells^22^. *CXCR4,* encoding an important chemokine receptor for PC homing and survival, was also differentially expressed only in IgG+ PCs from inflamed regions, consistent with prior findings in gut PCs from HIV and Crohn’s Disease^23^ (Extended Data Fig. 5a,d). Non-negative matrix factorization (NMF) of the scRNA-seq profiles of all 14 donors identified 10 programs (latent factors) underlying the variation across the cells (**Methods**), which we annotated based on high loading genes (Extended Data Fig. 6a, **Supplementary Tables 2, 3**). The programs for cell division (Program 1), cellular biosynthetic processes and oxidative stress (3), UPR response (4) and MHC-II-expression (7) increased in cells from inflamed regions (Fig. 2g, Extended Data Fig. 6b).

Multiple genes associated with inflammatory bowel disease (IBD) or UC through GWAS^23,24,25,26^ are highly expressed in IgG+ PCs from UC inflamed tissue, including genes related to oxidative stress (*GPX1, PRDX5, PARK7*)^15, 16^ and ER processes (*KDELR2*, *SDF2L1*)^27, 28^ (Extended Data Fig. 7a, b, **Methods**, pseudobulk DE analysis, FDR < 0.05). SDF2L1 is upregulated during the acute UPR and functions as an ER chaperone to facilitate protein cargo secretion^29^. Overall, genes associated with elevated antibody secretion, surface BCR expression, antigen presentation and cell division are upregulated in PCs from inflamed colon areas.

### PC expression profiles follow inflammation status rather than clonal membership

Examining the expression profiles of all cells from 6 representative PC clones from 3 donors with both inflamed and less inflamed colon areas (Fig. 3a,b, **Supplementary Table 4**), we found that PCs of the same clone separated mostly based on the inflammation status of their corresponding sample type. In all subjects with inflamed/less inflamed samples, the pairwise cosine distances of PC expression profiles was significantly larger between clonal members across inflammation states compared to clonal members within inflamed or less inflamed regions (Fig. 3c, one tailed t-test, P-value <= 10^-4^, Methods). With one exception (UC10), this was the case for each individual subject with matched inflamed and less inflamed samples (Extended Data Fig. 7c, one tailed t-test, P-val <= 10^-4^, **Methods**). Thus, expression differences between PCs from inflamed and less inflamed areas of the colon are apparent even for members of the same PC clone, suggesting that the local tissue environment is a dominant factor impacting their expression profiles.

**Figure 3:**
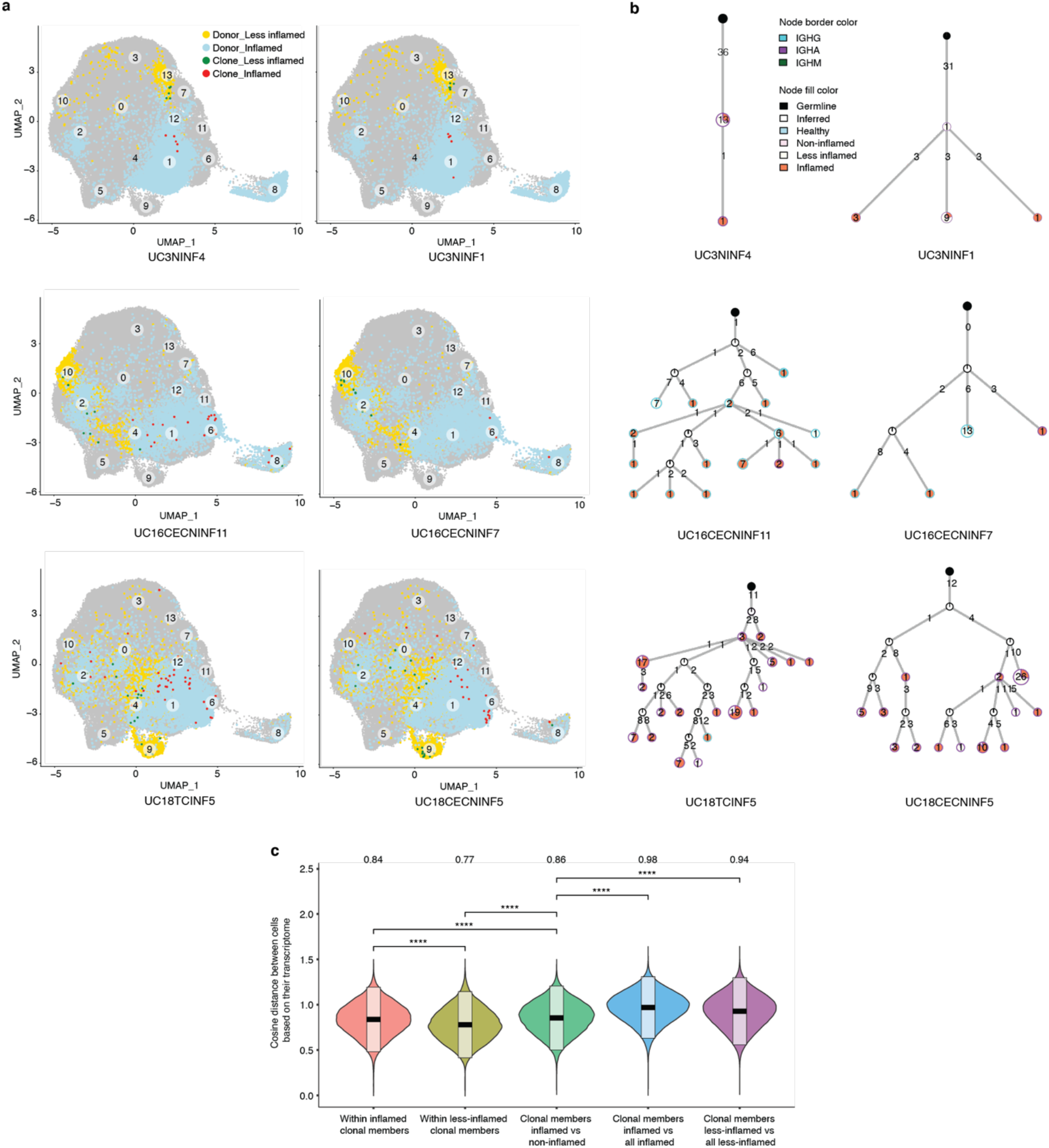
Transcriptomics of selected PC clones. **a)** UMAP plots highlighting clonal members of six selected PC clones from three different subjects as indicated (**Supplementary Table 4**). PCs from inflamed and less inflamed colon areas from the respective subject are highlighted in blue and yellow, respectively. Red dots indicate clonal members belonging to the selected clone from inflamed colon areas and green dots from less inflamed colon areas. **b)** Phylogenetic trees summarize the clonal relationship of all members within the selected clones. Trees are rooted on a theoretical germline member (black node), uncolored nodes indicate inferred intermediates and yellow and orange node colors indicate clonal members from less inflamed or inflamed colon areas, respectively. Node borders indicate the mAb isotype of the majority of cells in each node and the number in the center of each node indicates the number of cells represented in each node. Numbers on the connecting lines indicate the number of heavy chain mutations separating two nodes. **c)** Comparison of the pairwise cosine distances between PCs based on their transcriptome. Groups from left to right display the PC pair distance distributions between i) inflamed clone members, ii) less-inflamed clone members iii) inflamed and less-inflamed clone members iv) inflamed clone members and 100 randomly selected PCs of the donor that are inflamed and are not clone members v) clone members of less-inflamed samples and 100 randomly selected PCs of the donor that are less-inflamed and are not clone members. There is a significant difference between transcriptional distances between clone members based on their inflammation type.

### Antibodies from PCs in inflamed colon regions are not polyreactive or autoreactive

Polyreactivity, defined as non-specific binding to unrelated antigens, is selected against throughout B cell development^30^, but can arise during antibody affinity maturation^31^. The level of antibody polyreactivity in the human colon PC repertoire is unknown. Mouse derived antibodies from small intestine PCs reportedly have high levels of polyreactivity^32^, whereas antibodies isolated from PCs in the small intestine from both HCs and inflamed regions in Crohn’s disease patients display lower levels of polyreactivity^33, 34^. Similarly, autoreactivity is selected against during B cell development but can be generated during affinity maturation in germinal center reactions^31^ and is increased in peripheral B cell compartments of systemic lupus erythematosus patients^30^.

To evaluate polyreactivity and autoreactivity in mAbs from colon PCs in UC patients, we selected 152 mAbs from expanded clones from 5 UC patients and 2 HCs (Fig. 4a, **Supplementary Table 4**), and produced them as IgG1 mAbs irrespective of their original isotype in order to directly compare Fab reactivity. We defined polyreactivity as reactivity against at least two of the following antigens: single stranded DNA (ssDNA), double stranded DNA (dsDNA), lipopolysaccharide (LPS), insulin (INS) or cardiolipin (CAR)^31, 34^ (Fig. 4b). We defined autoreactivity as binding against HEP-2 cells (Fig. 4c). We detected polyreactivity only in 6 of 89 (6.7%) antibodies from inflamed colon areas, 2 of 39 (5%) antibodies from less inflamed colon areas and 0 of 15 (0%) antibodies from HCs; the frequency of polyreactive antibodies was not significantly different between PCs from inflamed colon compared to HCs (*p*=0.3 χ^2^ test) (Fig. 4b, Extended Data Fig. 8). Polyreactivity rates were lower than those in mAbs isolated from HC human memory B cells^31^ (overall 22.7%, *p*<0.005, χ^2^ test) or in mouse colon-derived antibodies^32^ (29%, *p*<0.005, χ^2^ test). Consistent with the low polyreactivity, only 1 of 152 mAbs (UC10NINF9) showed reactivity against HEP-2 cells (Fig. 4c).

**Figure 4:**
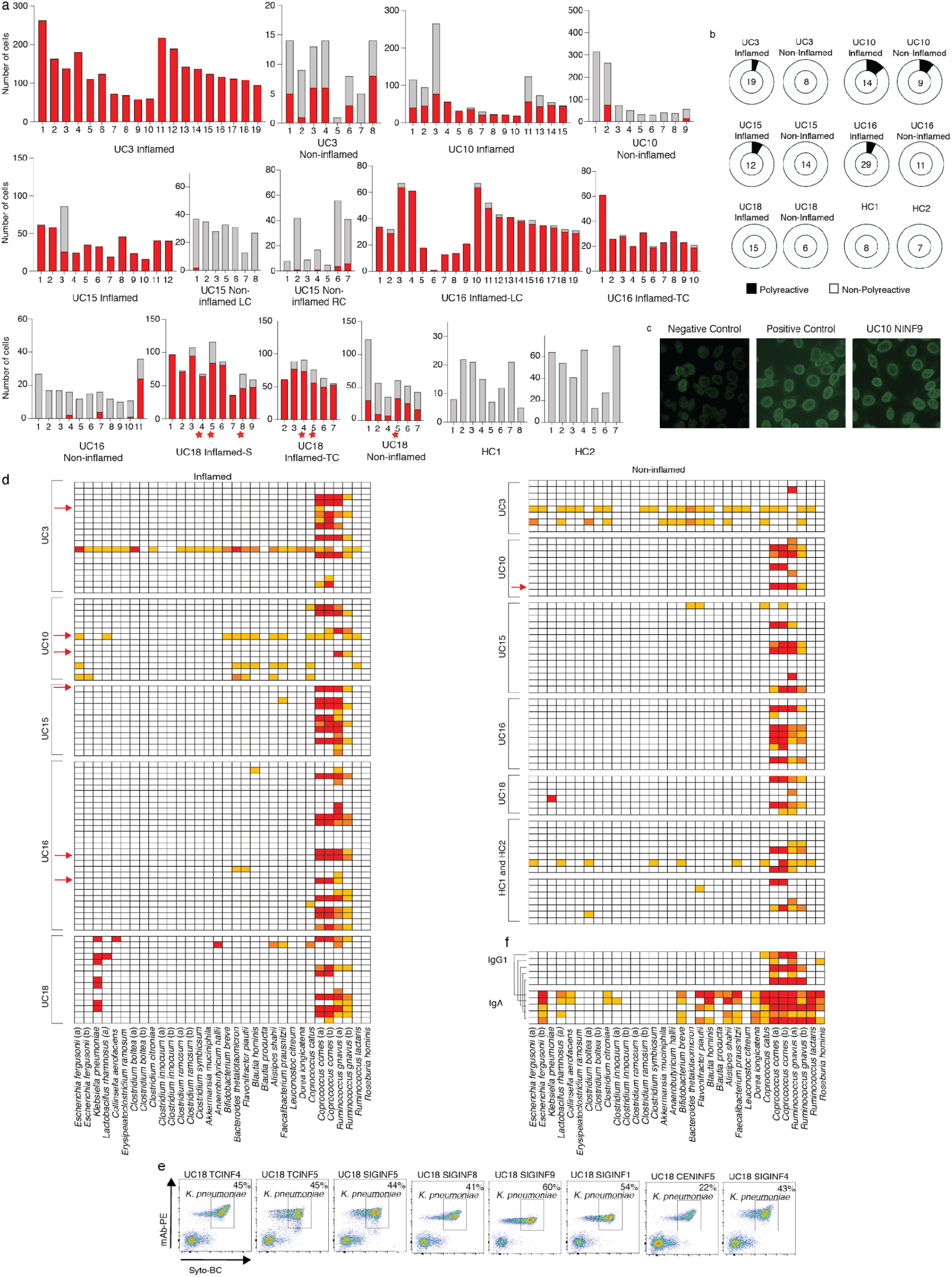
Functional testing of selected mAbs. **a,** Bar plots showing the clone size (number of cells, y-axis) of each selected and tested mAb (x-axis, **Supplementary Table 4**) as well as its expansion in inflamed (red bar) or less inflamed (grey bar) colon areas. mAbs are grouped based on the donor and colon area they were isolated from. The numbers below each clone correspond to the antibody names in Supplementary Table 4. LC=left colon, RC=right colon, TC=transverse colon, S=sigmoid colon, HC=healthy control. Red stars indicate clones from which *K. Pneumoniae* binding mAbs were isolated (see below). **b**, Pie charts summarizing the polyreactivity of all selected and tested mAbs. The number in the center of each pie indicates the number of mAbs tested and mAbs are grouped based on the donor and if they were isolated from an inflamed or less inflamed colon area (Extended Data Fig. 8). **c**, Representative HEp-2 cell IFA staining patterns of positive control and negative control serum as well as reactive mAb UC10NINF9 are shown. **d**, Heatmap showing the binding of all 152 selected mAbs against a panel of 32 bacterial strains in FACS. Each row represents one mAb and mAbs are sorted based on the donor and inflammation status of the colon region they were isolated from as indicated. White=<1% reactivity, light orange=1-5% binding, dark orange=5.1-10% binding and red=>10%binding. Red arrows indicate polyreactive mAbs (see above). **e**, representative FACS plots show the binding of reactive mAbs against bacterial strains used in (**d**). Antibody binding was detected using a mouse anti-human IgG antibody coupled to PE and bacteria were stained with SYTO-BC to exclude dead bacteria (**Methods**). **f**, heat map showing the binding of unrelated mAbs BG10-5, BG10-12, BG10-8, BG1-26, BG1-27^43^ in their IgG1 form and IgA1 or IgA2 form as indicated against a panel of 32 bacterial strains in FACS. Color coding as in (**d)**.

### Colon PCs produce mAbs with strong binding to some bacterial strains

Sequencing of IgA-covered microbiota revealed preferential targeting of colitogenic bacteria^6^ and PCs isolated from the small intestine of healthy subjects and Crohn’s disease patients show both specific and cross-species reactivity^34^. However, evaluating specific antibody-microbiota interactions is technically challenging because of the differences between stool samples, due to non-specific interactions such as polyreactivity and binding to B cell superantigens^35^.

We tested each of the 152 mAbs for their binding to single strains of a 32-strain bacterial panel that we assembled based on common members of the human microbiome and strains enriched in the microbiome of IBD patients^36–41^ (Fig. 4d). Consistent with prior reports of *VH3*-encoded heavy chains binding to a superantigen on *Coprococcus comes* as well as *Ruminococcus gnavus*^35^, 54 of 57 *VH3*-encoded antibodies showed strong binding against at least one of these strains (Fig. 4d) and 54 of 60 antibodies with strong binding carried a *VH3* heavy chain. Cross-strain reactivity with strong binding (>10% binding in FACS) to two or more bacterial strains (not counting non-specific superantigen binding as described above) was only detected in 4 of 152 antibodies (Fig. 4d), none of which were polyreactive. Aside from *R. Gnavus* and *C. Comes* binding, 5 of 152 antibodies showed strong binding to a single bacterial strain, *Klebsiella pneumoniae* (Fig. 4d, e). All *K. pneumoniae*-binding mAbs were isolated from one UC patient (UC18, Fig. 4d, e, Extended Data Table 1) and each of these mAbs represents an expanded PC clone that spans multiple colon regions with preferential expansion in inflamed regions (Fig. 4a, **red stars**).

To test if our 152 mAbs bind to stool bacteria, we grouped them into 12 mAb pools, including pools of *VH3* and non-*VH3* mAbs from inflamed, non-inflamed and HC colon-derived PCs, as well as one mAb pool with all polyreactive mAbs, (**Supplementary Table 5**), and tested for binding against stool from C57BL/6J and RAG-1-deficient mice (to avoid cross-detection of IgG-bound bacteria in human stool samples). mAB binding frequencies were low (0.3%-4.1%, Extended Data Fig. 9a), consistent with the observed low level of polyreactivity and cross-strain reactivity (above).

Further testing of the 152 mAbs against an array of 50 bacterial lysates^42^ (**Methods, Supplementary Table 6**), showed that 6 of 8 mAbs binding to *K. pneumoniae* in FACS also bound *K. pneumoniae* lysates, two of them with at least 8-fold lower cross reactivity against *Enterococcus faecalis* extract, *Bacteroides thetaiotaomicron* extract and *Bacteroides caccae* extract (Extended Data Fig. 9b, **Supplementary Table 6**). Another antibody from UC inflamed tissue bound *B. caccae* extract, two bound elongation factor Tu from *Bacteroides fragilis* and one bound *Prevotella intermedia* extract (Extended Data Fig. 9b, **Supplementary Table 6**). None of these binding mAbs were polyreactive (Extended Data Fig. 8, 9b). Finally, none of the 152 antibodies strongly bound common enteric viruses, including cytomegalovirus (CMV), Epstein-Barr virus (EBV) and rotavirus in ELISA (Extended Data Fig. 8). We conclude that mAb-microbiota interactions from local PCs include specific interactions with *K. pneumoniae* as well as non-specific superantigen interactions with *C. comes* and *R. gnavus* and that broad cross-strain reactivity is rare.

### IgA binds non-specifically to several microbiota

A significant fraction of microbiota in stool from mice and humans is bound by endogenously produced IgA^6^, but how much of that binding activity reflects specific Fab interactions with microbial surface antigens vs. non-specific binding is unknown.

To test if isotype switching changes the binding activity of antibodies to microbial targets, we chose two ‘orthogonal’ mAbs with known specificity to SARS-CoV-2^43^ and tested their IgG1 and IgA1 forms for binding to stool from C57BL/6J and RAG-1-deficient mice (Extended Data Fig. 9c). While IgG1 forms of both mAbs showed no binding, the same antibodies expressed as IgA1 showed significant binding to all stool samples tested, ranging from 4.9-6.9% (Extended Data Fig. 9c). To test if increased binding of microbiota to IgA1 mAbs is observed across different bacterial strains, we tested the same two antibody pairs as well as three additional SARS-CoV-2-specific IgG1/IgA1 pairs against our panel of 32 bacterial strains. While we detected minimal binding of IgG1 mAbs (Fig. 4f), 15 out of the 34 strains that were neither *R. gnavus* nor *C. comes* were bound by at least two of the IgA1 mAbs. We conclude that a significant fraction of microbiota show non-specific binding to unrelated IgA1 molecules and that those interactions likely involve residues outside of the antigen binding site.

## DISCUSSION

Immune responses to the microbiome play an important role in UC^2^, and the protective role of FCGR2A^3, 4^, as well as expansion of PCs in UC crypt infiltrates^3^, places PCs in the center of investigating novel therapeutic avenues in UC. Our results suggest that the clonal landscape in the healthy colon is dominated by IgA1/IgA2 PCs, which overlap significantly between different colon areas, and that UC disrupts this landscape, mostly through IgG1 dominant clones that show limited overlap with IgA1/IgA2 clones which could be a result of limited class switching within germinal center reactions^44^. IgG1 dominant clones in inflamed regions also frequently contain multiple identical clonal members suggestive of a memory B cell origin in the setting of a secondary immune response. Interestingly, even UC patients in remission show an expansion, albeit less pronounced, of IgG1 PC clones that might be responses that remain dormant in remission, but may readily expand upon recurrence of inflammation. Future longitudinal studies of UC patients will help elucidate this.

Polyreactivity and autoreactivity are not significantly enriched in mAbs from inflamed colon areas and are overall rare in all disease states, a finding consistent with the highly specific mAb interactions with selected bacterial strains we identified. The expansive reactivity of multiple PC clones with *K. pneumoniae* in the inflamed samples from one subject raises the question if certain pathogenic bacterial strains are preferentially targeted in the setting of UC inflammation. Beyond specific interactions, we also found significant non-specific *VH-3* binding to *R. gnavus* and *C. comes,* and demonstrated that an isotype switch from IgG1 to IgA1 in unrelated SARS CoV-2-specific mAbs can lead to nonspecific binding against selected bacterial strains. These findings underscore the multitude of mAb-microbiome interactions and the importance of investigations at the monoclonal level to understand the role of specific B cell responses in UC. Certainly, the upregulation of MHCII-related genes in PCs from inflamed areas suggests that such specific B cell responses could also impact inflammatory T cell responses. Further studies of these interactions and the antigens involved are needed.

## Methods

### Human Subjects

All work with human samples was performed in accordance with approved Institutional Review Board (IRB) protocols which were reviewed by the IRB at Massachusetts General Hospital (MGH), Boston. HCs as well as UC patients were recruited through a patient cohort that has been created at MGH under IRB protocol 2004P001067, “Prospective Registry in Inflammatory Bowel Disease Study at Massachusetts General Hospital”. HCs were overall healthy individuals without any history of IBD, other autoimmune disease, infectious colitis or colon cancer. UC patients were included based on carrying a clinical diagnosis of ulcerative colitis. UC patients were determined to either have active disease or to be in remission based on macroscopic and histopathologic evaluation of the mucosa in the setting of endoscopic evaluation or of a resection sample (Extended Data Table 1). None of the subjects had any known history of infectious colitis. Resection samples were obtained in the setting of total colectomies.

### Single cell dissociation from fresh colon mucosa samples

Biopsy bites or mucosal resection samples were immediately processed and mucosal samples placed into cryovials containing Advanced DMEM F-12 media (ThermoFisher Scientific) and placed on ice for transport. Single-cell suspensions from collected mucosal samples were obtained using a modified version of a previously published protocol^47^ as detailed below. Mucosal samples were first rinsed in 30 mL of ice-cold PBS. Each individual sample was then transferred to 5mL of enzymatic digestion mix (Base: RPMI1640, 100 U/ml penicillin (ThermoFisher), 100 μg/mL streptomycin (ThermoFisher), 10 mM HEPES (ThermoFisher), 2% FCS (ThermoFisher), and 50 μg/mL gentamicin (ThermoFisher), freshly supplemented immediately before with 100 mg/mL of Liberase TM (Roche) and 100 μg/mL of DNase I (Roche), and incubated at 37°C with 120 rpm rotation for 30 minutes. After 30 minutes, the enzymatic dissociation of the lamina propria was quenched by addition of 1ml of 100% FCS (ThermoFisher) and 80 μL of 0.5M EDTA and placed on ice for five minutes. Samples were typically fully dissociated at this step and after gentle trituration with a P1000 pipette filtered through a 40mM cell strainer into a new 50 mL conical tube and rinsed with PBS to 30 mL total volume. This tube was spun down at 700g for 10 minutes and resuspended in 500ul of PBS with 5% fetal bovine serum (FBS). Cell suspensions were then stained with anti-human CD27 PE (BD), CD38 FITC (BD) and CD19 APC (BD) and incubated for 20 minutes at 4°C before they were washed and resuspended in PBS with 5% FBS. PCs were then sorted using a Sony MA900 cell sorter by gating on live cells in the forward scatter and side scatter and on CD38-FITC and CD27-PE double-positive cells (Fig. 1b). After sorting, cells were washed and counted using a hemocytometer and microscopy, before resuspending up to 10,000 cells in a volume of 32μl for 5’ single-cell RNA-Seq (see below).

### 5’ scRNA-seq library generation

Cells were separated into droplet emulsions using the Chromium Next GEM Single-cell 5′ Solution (v1.1) and the 10x Chromium Controller. 5,000-10,000 cells were loaded per channel of the Chromium Next GEM single-cell 5’ (v1.1) Chip G. Following cell lysis, barcoded mRNA reverse transcription, and cDNA amplification, a 0.6X SPRI cleanup was performed, and the supernatant was set aside for Feature Barcoding library construction as instructed by the Chromium NextGEM single-cell V(D)J v1.1 protocol. A final elution of 45μL was saved for further library construction. Using the saved supernatant of the 0.6X cDNA cleanup, Feature Barcoding libraries were completed according to the 5’ Next GEM (v1.1) Feature Barcoding library construction methods provided by 10x Genomics. Gene expression and V(D)J libraries were created according to the manufacturer’s instruction (10x Genomics), which includes enzymatic fragmentation, adaptor ligation, and sample index barcoding steps. The V(D)J libraries were created from the original following the Chromium NextGEM single-cell V(D)J v1.1 protocol.

### scRNA-seq library sequencing

Gene expression, feature barcoding libraries, and BCR enriched V(D)J libraries were sequenced either on a Nextseq500 (Illumina) using a high output 150 cycle flowcell, with the read configuration Read 1: 28 cycles, Read 2: 96 cycles, Index read 1:8 cycles or on a HiSeq X (Illumina), using a 150 cycle flowcell with the read configuration: Read 1: 28 cycles, Read 2: 96 cycles, Index read 1: 8 cycles. Feature Barcoding libraries were spiked into the gene expression libraries at 10-20% of the sample pool prior to sequencing. All BCR enriched V(D)J libraries were pooled together and sequenced on a NextSeq500 (Illumina) using the same parameters as previously mentioned.

### mAb production

mAb VDJ heavy chain and VJ light chain sequences of selected mAbs were produced as minigenes and cloned into IgG1 heavy chain, kappa light chain or lambda light chain expression vectors as previously described^30^. After cloning into expression vectors, matching mAb heavy chain and light chain plasmids were co-transfected into Expi293F cells following the manufacturer’s instructions. Briefly, heavy chain plasmid DNA and light chain plasmid DNA were diluted in 1.5ml Opti-Plex Complexation Buffer (Invitrogen) before mixing with 80µl ExpiFectamine 293 Reagent (Invitrogen) diluted in 1.4 ml Opti-Plex Complexation Buffer (Invitrogen). Mixture was incubated for 15 minutes at room temperature before adding to 25 ml of Expi293F cells at a density of 3.0 × 10^6^ viable cells/ml and incubation in a shaker incubator according to the manufacturer’s instructions. ExpiFectamine 293 Transfection Enhancer 1 and 2 (150 µl and 1.5 ml, respectively) were added 18 hours post-transfection. mAb containing supernatants were harvested after 7 days by centrifugation of cells at 3,000g for 20 minutes and transfer of supernatants into 50 ml Falcon tubes (Fisher Scientific). IgGs were purified from transfection supernatants using protein G coupled agarose beads (GE) according to the manufacturer’s instructions.

### mAb ELISA testing

mAb concentrations were determined measuring absorbance at 280 nm using Nanodrop 2000c (ThermoFisher) or IgG specific ELISA as previously described^48^. mAb reactivities to EBV, CMV and rotavirus were determined using IgG detection kits against the respective viruses: EBV and CMV (abcam), rotavirus (amsbio) and the concentration of mAb used was 1μg/ml. Polyreactivity ELISAs against antigens ssDNA, dsDNA, LPS and insulin and ELISA against cardiolipin (Sigma) were performed as previously described^48, 49^. Similar to prior studies, polyreactivity was defined as reactivity to two or more antigens including single stranded DNA, double stranded DNA, lipopolysaccharide (LPS), insulin or cardiolipin^34, 48^. As previously described, mAbs ED38, JB40 and mGO53^30^ were used as strongly polyreactive, intermediate polyreactive and non-polyreactive control mAbs, respectively.

#### Protein microarray and detection

Bacterial proteins and extracts were arrayed in quadruplicate on 16 pad nitrocellulose slides (Maine Manufacturing) at a concentration of 0.2mg/ml using a Spotbot Personal Microarrayer (Arrayit). Recombinant proteins and bacterial extracts were diluted from stocks to 10mM Tris (pH7.4), 20% glycerol, and 0.1% SDS. To probe the microarray, pads were blocked for 1 hour in SuperBlock (ThermoFisher) and then probed with human monoclonal antibodies diluted 1 to 250 (final concentration 1.2μg/ml) in SuperBlock for 1 hour. The pads were washed three times in PBS with 0.05% Tween 20; then, anti-human IgG secondary antibody labeled with Dylight 650 (Invitrogen) was applied at 1 to 1000 (∼0.5μg/ml) for 1 hour in SuperBlock. The pads were then washed three times with PBS-tween, distilled water, air-dried and scanned using a GenePix 4000B imager (Axion).

### Bacterial isolation and verification via 16S sequencing

All bacterial strains used in this study are detailed in Fig. 4d. Bacteria were cultured and stored using the following protocol: bacteria were struck out and streaked from frozen bacteria stocks onto Cullen-Haiser Gut (CHG) or yeast casitone fatty acid agar with carbohydrates (YCFAC) media plates under anaerobic conditions. Plates were incubated at 37°C, under anaerobic conditions for 24-72h, until isolated bacterial colonies were obtained. Next, a single bacterial colony was inoculated into a new 15mL falcon tubes (Corning) containing CHG or YCFAC media. Liquid cultures were incubated at 37°C, under anaerobic conditions, for 24-72h, until optical density at 600nm (OD600) of 0.2 or higher were obtained. Liquid cultures were combined with 50% glycerol, 1X PBS solution (1:1 ratio) and aliquoted into cryovials. Cultured strains were verified via 16S rRNA v1-v9 sequencing and further aliquots were stored at −80°C for flow cytometry analysis at a later time. Verification of cultured strains via 16S sequencing was conducted as follows: 5μL harvested bacterial culture was aliquoted to a corresponding well in a sterile 96-well PCR plate (ThermoFisher) and added 50µL of hot-shot lysis buffer was added to each well containing bacterial aliquots, as well as three additional, empty reagent control wells. Plates were sealed and placed into PCR thermocycler (Bio-Rad) for 10 min at 95°C, followed by a holding stage at 12°C. 50μL neutralization buffer was added to each well, mixed gently and stored plates at −20°C. Plates with template DNA were thawed on ice and PCR reagents (1.5µL 10mM 16S rRNA forward prime (IDT), 1.5µL 10mM 16S rRNA reverse primer (IDT), 12.5µL OneTaq 2X MasterMix (NEB) and 5µL nuclease-free water) were added to new sterile 96-well PCR plate and mixed before addition of template DNA. Three wells of PCR reagents without template DNA were added as reagent controls. Plates were sealed and placed into PCR thermocycler for the following protocol: step 1, 94°C for 30 s; step 2, 94°C for 30 s; step 3, 55°C for 60 s; step 4, 68°C, 90 s (cycle steps 2 to 4, 30 times); step 5, 68°C for 5 min; step 6, 12°C, hold. Plates were then submitted to Genewiz Inc. for enzymatic purification and 16S V1-V9 Sanger sequencing, with special instructions to use 2.5µL of 10mM 16S rRNA forward primer for sequencing. The taxonomic identity of each strain was verified by importing 16S rRNA data into Geneious Prime 2021.2.2 and performing standard nucleotide BLAST via blastn suite software (NIH, National Library of Medicine, https://blast.ncbi.nlm.nih.gov/Blast.cgi).

### Bacterial prep for flow cytometry

Verified bacteria were washed, diluted, and stained for downstream flow cytometry analyses as follows: bacteria were thawed on ice, resuspended, filtered through a sterile 50µm CellTrics filter (Sysmex America), transferred to a new 1.5-mL tube, and centrifuged at 8000 g for 5 min. Supernatants were removed by aspiration, and pellets resuspended in 1.5 mL cold PBS 0.25% BSA (Sigma), and centrifuged at 8000 g for 5 min. Supernatants were removed by aspiration and pellets resuspended in 1.0 mL cold PBS 0.25% BSA. Optical densities were recorded at 600 nm (OD_600_) via NanoDrop 2000c (ThermoFisher) and 2.0 mL cuvette (Fisherbrand) to estimate bacterial culture densities. Suspensions were adjusted to OD_600_ of 0.1 to 0.2, 50 μL/well was aliquoted to a Nunc 96-well, V-bottom, polypropylene plate (ThermoFisher). Next, mAbs were diluted to a concentration of 2.0 µg/mL in cold PBS 0.25% BSA in a new 96-well plate. 50 µL of diluted mAbs were transferred to the 96-well plate containing 50 µL/well of bacterial suspension (final working mAb concentration is 1 µg/mL). mAb testing was performed for binding to bacteria as described below.

### Mouse stool sample collection and homogenization

Fresh fecal pellets were collected from adult WT C57BL/6J (Jackson Laboratory, Stock #000664) and Rag1^tm1Mom^ (Jackson Laboratory, Stock #002216) mice, pooled by genotype, and stored at −80°C. Mouse fecal pellets were thawed and resuspended in 10 mL of BIOME-preserve anaerobic medium, were transferred to a gentleMACS C tube (Miltenyi), and homogenized via gentleMACS Dissociator (Miltenyi) for 3 cycles of 61 s on the ‘intestine’ setting. All stool homogenates were aliquoted into 2.0-mL cryovials (VWR) and stored at −80°C.

### Stool sample preparation for flow cytometry

Homogenized stool samples were washed, diluted, and stained for downstream flow cytometry and FACS analyses as follows: stool samples were thawed on ice, resuspended, and transferred to a new 1.5-mL tube. Samples were centrifuged at 50 g for 20 min to pellet large debris. Supernatants were filtered through a sterile 50µm CellTrics filter (Sysmex America) and transferred to a new 1.5mL tube. Samples were centrifuged at 8000 g for 5 min and supernatants removed by aspiration. Pellets were resuspended in 1.0 mL cold PBS 0.25% BSA. Optical densities were measured at 600 nm (OD_600_) via NanoDrop 2000C and 2.0 mL cuvette to estimate bacterial culture densities. Suspensions were adjusted to OD_600_ of 0.1 to 0.2 and 50 µL/well added to a Nunc 96-well V-bottom plate. mAbs were diluted to a concentration of 20.0 µg/mL in cold PBS 0.25% BSA in a new 96-well plate. 50 µL of diluted mAbs were transferred to the 96-well plate containing 50 µL/well of bacterial suspension (final working mAb concentration is therefore 10 µg/mL). mAb testing was performed for binding to stool samples as described below.

### mAb testing for binding to bacteria and stool samples

We continued from aforementioned prep steps as follows: after addition of mAbs plates were covered with foil Microseal (Bio-Rad), incubated on ice for 30 min, and centrifuged at 5000 g for 10 min. Cover was removed and supernatant decanted. Pellets were resuspended in 250 µL cold PBS 0.25% BSA and centrifuged at 5000 g for 10 min after which supernatant was decanted. Pellets were resuspended in 50 µL cold PBS 0.25% BSA with PE-conjugated mouse anti-human IgG (1:100, BD Biosciences), or PE-conjugated goat-anti human IgA (1:800), incubated on ice for 20 min, and centrifuged at 300 g for 1 min. After that, 50 µL/well of cold PBS 0.25% BSA with SYTO BC nucleic acid stain (1:500, ThermoFisher) was added. After incubation at 10 minutes on ice, the plate was centrifuged at 5000 g for 10 min and the supernatant decanted. Pellets were resuspended in 250 µL cold PBS 0.25% BSA and centrifuged at 5000 g for 10 min, supernatant removed and pellets resuspended pellets in 100 µL cold PBS 0.25% BSA. Flow cytometric analyses were conducted via CytoFLEX S flow cytometers (Beckman Coulter), visualizing intact bacteria by gating on FSC^+^SSC^+^FITC^+^ cells, and visualizing mAb-bound bacteria by gating on intact bacteria PE^+^ cells.

## QUANTIFICATION AND STATISTICAL ANALYSIS

### Preprocessing of scRNA-seq and V(D)J readouts

mRNA and VDJ sequence reads were mapped to the reference human genome GRCh38-3.0.0 with the cloud-based Cumulus workflows^50^, using the CellRanger 3.0.2 software pipeline.

### scRNA-seq analysis

For the mRNA data integration, count normalization, dimensionality reduction, clustering, cell scoring, and cluster marker genes detection Seurat R package^45, 46^ was employed.

#### Preprocessing and Batch correction

Cells which either do not have 10x standard high-quality heavy and light chain V(D)J sequences, or have more than 10% of their transcriptome reads coming from mitochondrial genes were filtered out before the downstream transcriptome analysis. For the UMI count normalization step, gene expression counts for each cell were divided by the total counts for that cell and multiplied by 10^6^, which was then log-transformed using log1p. Top 2000 variable genes were identified with utilization of variance stabilizing transformation (vst), where a local polynomial regression (loess) model was fit to model relationship of log(variance) and log(mean) and gene values were standardized using the observed mean and the expected variance given by the fitted line. Gene variance was then calculated on the standardized values after clipping to a maximum, which was defined as the square root of the number of cells. Batch effects, defined as the batch of the sequencing run, were regressed out from the normalized count values with the ComBat algorithm^51^ implemented in SVA R Package version 3.38.0. Finally, the resulting residuals were scaled and centered.

#### Dimensionality reduction, graph clustering, and UMAP visualization

Dimensionality reduction was done with PCA identifying the first 50 principal components. For clustering of the cells into expression clusters, a *k*-nearest neighbor (*k*NN) graph of the cells was constructed (k=20) using the 50 principal components. Next, this kNN graph was used to generate the shared nearest neighbor (sNN) graph by calculating a Jaccard index between every cell and its *k* nearest neighbors. Then the Leiden algorithm^52^ was used to find the clusters of the cells based on the generated sNN graph, with a resolution of 0.024 decided based on the identifiability of the marker genes. Expression levels of immunoglobulin genes were discarded during the clustering step. Uniform manifold approximation and projection (UMAP)^53^ algorithm was run on the first 50 principal factors to obtain the 2D projections of the cells.

#### Scoring gene signatures

Module scores of the cell cycle genes and the antigen presentation genes were calculated as previously defined in [47], where for each gene in the module gene-set, 100 genes were randomly selected as control genes.

#### Differential expression analysis

Differentially expressed genes between UC inflamed and healthy samples were identified with pseudobulk differential expression analysis using the DESeq2 R package^54^, where counts of each gene were aggregated at the sample level. Multiple testing correction was performed with the Benjamini-Hochberg procedure. Genes that had FDR < 0.05 were accepted as significantly differentially expressed between UC inflamed and healthy samples.

#### Non-negative matrix factorization

Non-negative matrix factorization of the integrated single cell count matrix was done by utilizing the Consensus Non-negative Matrix factorization (cNMF)^55^ Python package. Number of high variance genes that were used for running the factorization was set to 3000 and the loss function for NMF was *frobenius*. Optimal number of latent factors was decided based on the identifiability of the independent pathways verified by the gene set enrichment analysis. The gene s with significantly high loadings per factor were defined to have loadings greater than 3 interquartile ranges higher than the 75th percentile. Gene set enrichment of the selected genes was performed with the clusterProfiler R package^56^.

### Antibody repertoire analysis

#### Preprocessing and B cell clone calling

The V(D)J contig assembly algorithm from 10x Genomics (https://support.10xgenomics.com/single-cell-vdj/software/pipelines/latest/algorithms/assembly) takes many forms of noise specific to scRNA-seq data into account, while generating the assembled V(D)J sequences. Nevertheless, only the cells with high-quality heavy and light chain V(D)J contig sequences were selected and V(D)J gene annotations were assigned by using IGBLAST (version 1.14.0) software with the Change-O R package^57^. Cells with more than one high-quality heavy or light chain sequence (*i.e.* double expressors) were excluded from the downstream repertoire analysis. Donor-specific B cell clones were identified by utilization of the Change-O R package on the combined cell population of all collected biopsies of the donor. The appropriate threshold for trimming the hierarchical clustering into B cell clones was found by inspecting the bimodal distribution of the distance between each sequence in the data and its nearest-neighbor.

#### Comparison of isotype percentage between samples of different disease status

For isotype distribution comparison analysis, for each biopsy sample one representative member of each clone with unique heavy and light chain sequences was selected. Next, the percentage of the IgA1, IgA2, IgG1, IgG2, IgG3, IgG4 and IgM isotypes within the selected cells was calculated for each sample. Statistical significance of the differences between the distributions of the isotype percentages between samples with different disease status were tested with both non-parametric wilcoxon rank sum test and Dirichlet-multinomial regression, implemented as DirichReg function in DirichletReg R package^58^, to account for the fact that the percentage values of all isotypes within a sample sum up to 100.

#### Mutational load analysis

Replacement and silent mutation inference based on the scRNA-seq VDJ sequences of the donor cells^59^ was performed by the Shazam R Package^60^, where the region definition parameter was set to be “IMGT_V_BY_SEGMENTS” which provides no subdivisons and treats the entire V segment as a single region.

#### CDR3 region length and amino acid physicochemical property analysis

CDRH3 length was defined based on IMGT definition^61^ with the addition of two conserved amino acid residues to assist in clonal analysis^62^. CDRH3 amino acid charges were calculated by the Alakazam R package^57^ using the method of Moore^63^, excluding the N-terminus and C-terminus charges, and normalizing by the number of informative positions. Hydrophobicity scores were calculated with the Alakazam R package using the method of Kyte and Doolittle^64^. Shannon entropy values were calculated using the Alakazam R package. For each donor, the transcriptome cluster specific Hill diversity index^65^, improved by Chao et al^66, 67^ was calculated by setting the diversity order equal to 1 with Alakazam R package. For each run, the number of bootstrap realizations was set to be 1000, and the minimum number of observations to sample was set to 20.

#### Clonal overlap analysis

To compare the clonal repertoire between two samples, 100 cells were randomly sampled from each sample 1,000 times, and for each pair of random selections we calculated the percentage of cells that belong to clones that have members in each of the two random samples. The distribution of these 1,000 clonal overlaps is taken as a measure of repertoire similarity between the two compared samples. The same testing schema is repeated with one clonal representative cell per sample to control for the bias that might have been introduced due to the greater number of cells in the expanded clones.

#### Analysis of isotype conservation within clones across regions with different inflammation status

Dominant isotype conservation across inflammation status in the expanded clones (n > 9) that span both inflamed and less-inflamed regions of the donor was tested by calculating the Spearman’s correlation coefficient between the clonal percentage differences of the IgG and IgA cells coming from the inflamed samples (column 4 minus column 6 of Extended Data Fig. 1d) versus the clonal percentage differences of the IgG and IgA cells from the less-inflamed samples (column 3 minus column 5 of Extended Data Fig. 1d).

#### Comparison of transcriptional distance between cells based on the clonal membership

To test the partitioning of the clones on the expression landscape, for each clone which has members from both inflamed and less inflamed regions, the pairwise cosine distance was calculated between: i) members of the clone from samples with the same inflammation status, ii) members of the clone from samples with a different inflammation status, iii) 200 randomly selected cells of the donor that are not members of the clone from samples with the same inflammation status, iv) 200 randomly selected cells of the donor that are not members of the clone from samples with different inflammation status. The significance of the difference between the distributions of these cosine distances was tested with a one-tailed t-test.

## Acknowledgements

We thank Heather Kang for editorial assistance with the manuscript and figures. We thank all study participants who devoted time to our research and we thank the clinical staff and research coordinators lead by Helena Lau at MGH. We thank Patricia Rogers and Natan Pirete (Broad Institute) for support with cell sorting. Cell sorting was performed at the Flow Cytometry Facility of the Broad Institute, sequencing of pre-constructed DNA libraries was performed at the Genomics Platform (Broad Institute). This work was supported by NIH (RC2 DK114784 and P30 DK043351 to R.J.X.) and The Leona M. and Harry B. Helmsley Charitable Trust.

## Author contributions

J.F.S., B.E., D.B.G., A.R. and R.J.X. conceived the study, designed experiments and wrote the manuscript with contributions from J.D. and other authors, J.F.S., A.A. and A.T.J. established and orchestrated the patient cohort, J.F.S established and performed cell staining and sorting, B.E. performed computational analysis with guidance from A.R., B.E. and J.F.S. analyzed repertoire and transcriptome data with guidance from A.R., J.F.S and A.H. performed mAb cloning, production, purification and binding characterization. E.B., D.S. and H.V. grew bacterial stocks and assisted in mAb bacterial binding experiments. L.W.D. and C.O.E. developed and performed mAb binding assay to bacterial lysates, T.D., D.P. and A.R. developed and performed scRNA-seq library preparation and coordinated single cell RNA-seq.

## Additional information

Supplementary information is available for this paper.

## Supplemental Information

**Extended Data Figure 1:**
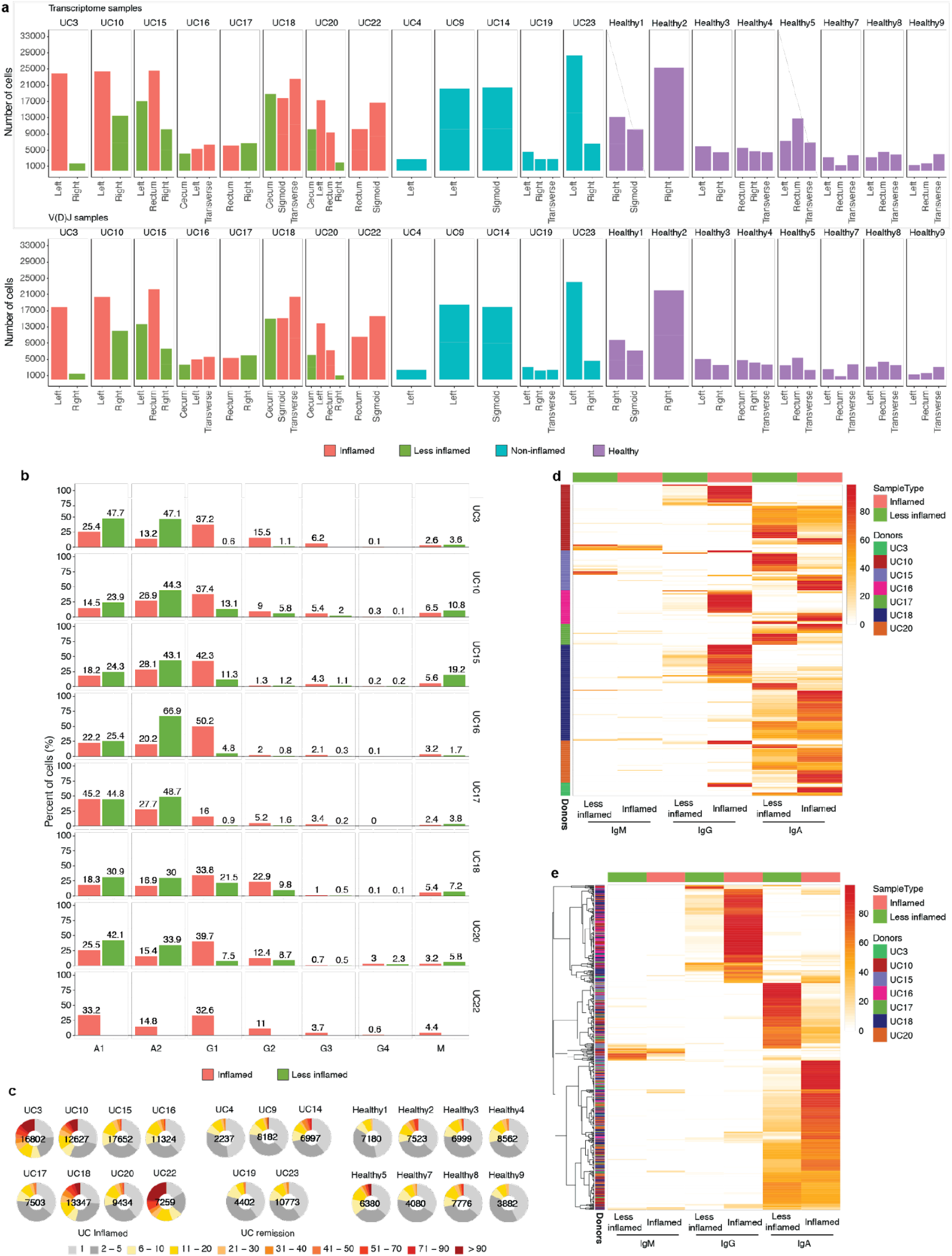
Isotype analysis and clonal expansion. **a,** Bar plots displaying the number of cells used in the downstream analysis (y-axis) from various colon regions (x-axis) of each subject. The upper graph shows the number of cells for which transcriptome profiling information was obtained and the lower graph shows the number of cells for which productive V(D)J sequence were obtained (**Methods**). **b,** Bar plots displaying the median percentage of each immunoglobulin isotype within the samples (y-axis) of each donor with at least one area of inflammation stratified by the sample inflammation status as indicated (x-axis). **c**, Pie charts show the expansion of differently sized PC clones for each donor among UC patients with inflammation (left), UC patients in remission (middle) and healthy controls (right). Numbers in the center of the pie charts represent the total number of cells analyzed in that particular plot. **d, e**, Heat maps showing isotypes of expanded clones that are shared between inflamed and less inflamed colon areas. The heat maps display the percentage of each immunoglobulin isotype – inflammation status pair within each expanded clone with more than 9 cells. The threshold of 10 cells was arbitrarily chosen to capture larger clones. Each row sums up to 100% and represents one expanded clone. Each column stands for the isotype-inflammation status pair. Heat maps are organized based on donor (**d**) or enrichment in isotype – inflammation status pair (**e**).

**Extended Data Figure 2:**
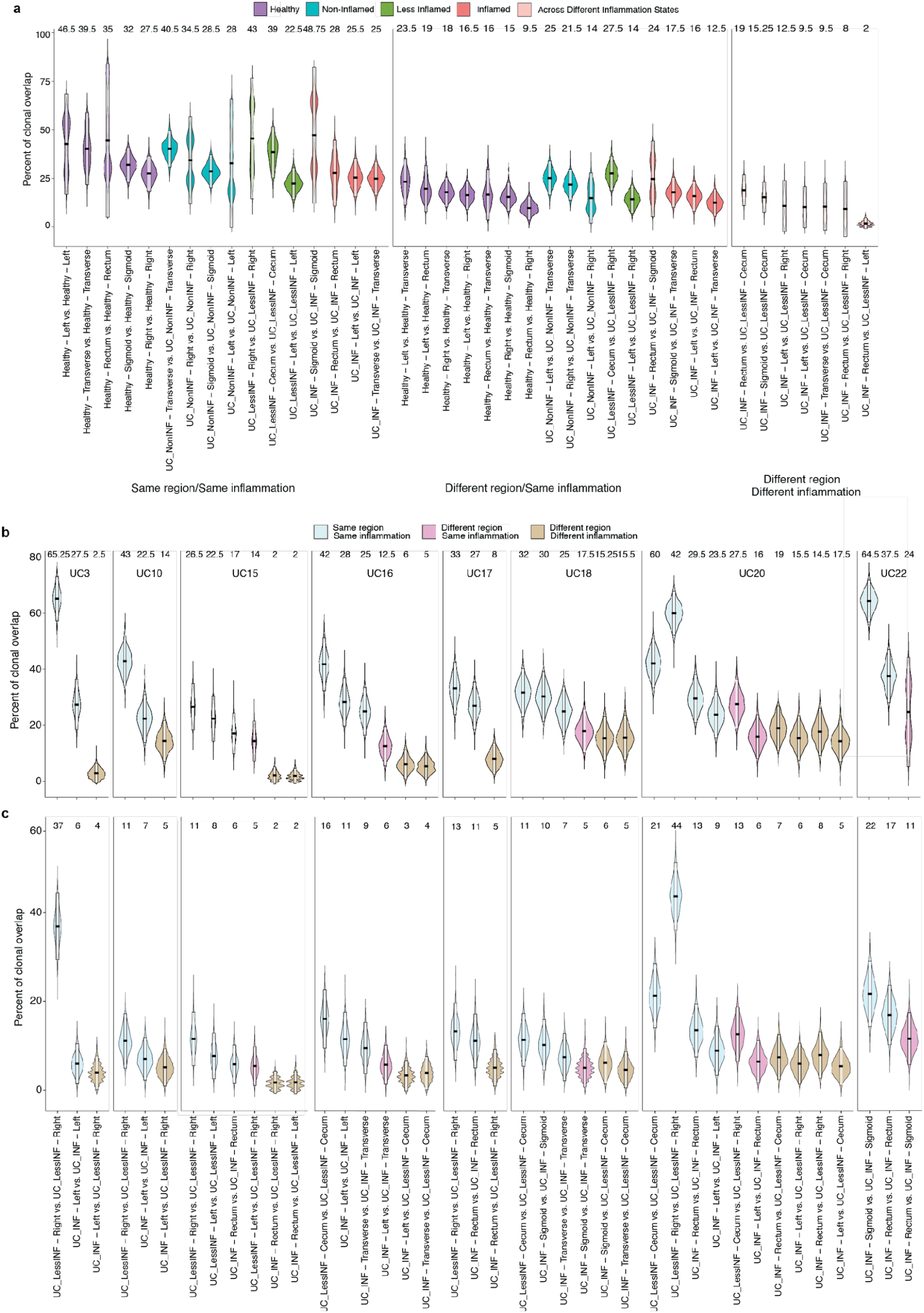
Clonal overlap between PCs from different colon regions. **a-b,** Violin plots display the distribution of the percentage of the shared clones between randomly sampled sets of PCs (**Methods**). In (**a**) the two random samples of a specific donor that are to be evaluated for clonal overlap can belong to i) same colon region (left), ii) different colon region with same inflammation status (center) and iii) different colon region with different inflammation status (right) as indicated. In contrast to Fig. 1f each colon region is analyzed separately. Purple violin plots represent HC samples, turquoise plots non-inflamed samples from subjects in remission, green plots less-inflamed and red plots inflamed samples in a UC patient with inflammation as indicated. Pink violins indicate samples that are being compared across different inflammation states. In (**b**) samples are separated by donor with blue violin plots representing overlap within the same colon region, pink different regions with the same inflammation status and beige different regions with different inflammation status. For x-axis labeling, see (**c**). **c**, Violin plots displaying the distribution of the percentage of the shared clones between randomly sampled sets of PCs (**Methods**). As opposed to Extended Data Fig. 2b, expanded clones in each sample are collapsed to one cell. The two random samples of a specific donor that are to be evaluated for clonal overlap can belong to the same tissue sample (blue), different colon region with same inflammation status (pink) and different colon region with different inflammation status (beige). The boxes represent −2 standard deviation (lower portion), mean (black line), and + 2 standard deviation (upper portion). The values above each violin plot represent the median values of the distribution.

**Extended Data Figure 3:**
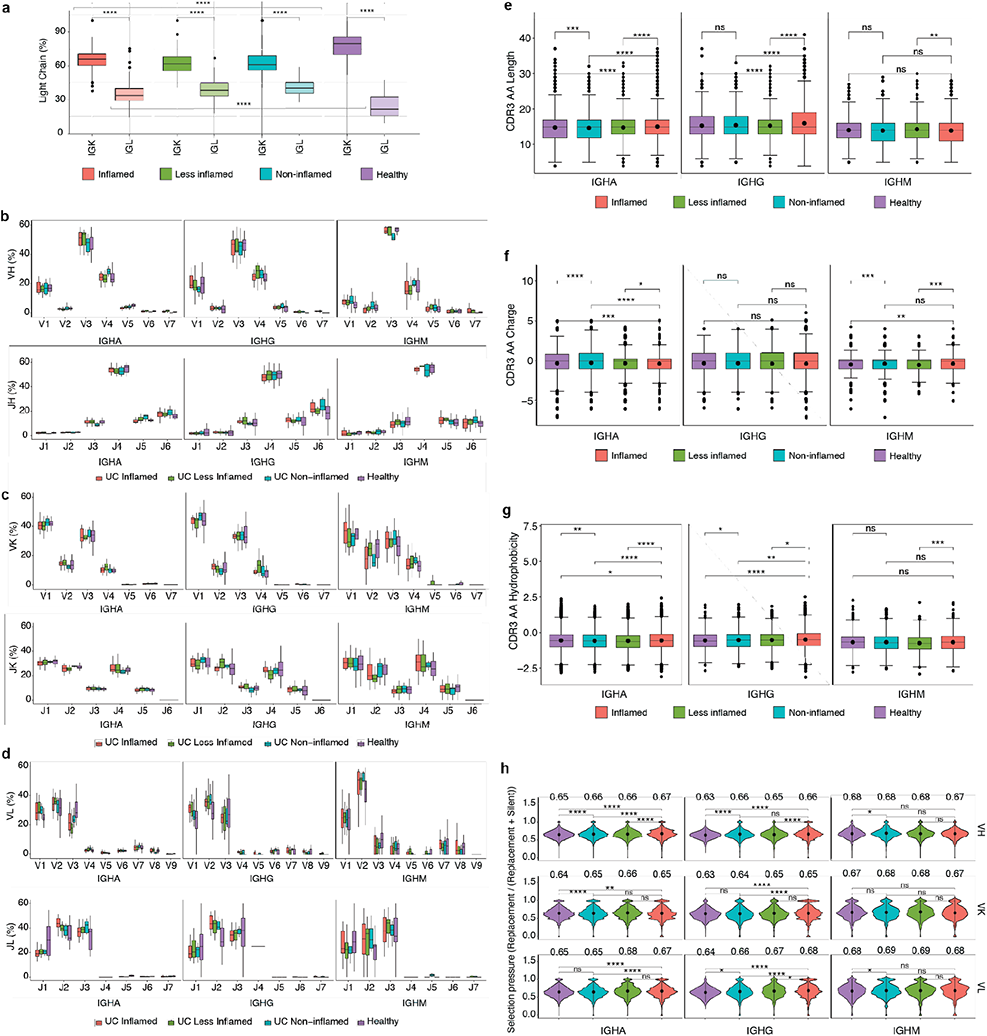
Colon PC antibody repertoire. Statistically significant differences are indicated with brackets. **a,** box plots display antibody light chain usage with kappa light chains shown in dark and lambda light chains in light colors. Antibodies are stratified by the disease state of the source tissue as indicated. Brackets indicate statistical significance using a one-sided non-parametric Wilcoxon rank-sum test \*\*\*\**P ≤* 0.0001. **b-d,** box plots showing the V- and J-gene usage of antibody heavy chains (**b**), kappa light chains (**c**) and lambda light chains (**d**) as indicated. Red plots represent antibodies derived from PCs in inflamed tissue, green plots antibodies from less inflamed tissue of UC patients with inflammation, turquoise plots antibodies from non-inflamed tissue in UC patients in remission and purple plots antibodies from HCs. **e-g,** box plots displaying the CDRH3 amino acid length (**e**), CDRH3 amino acid charge (**f**) and CDRH3 amino acid hydrophobicity (**g**) of antibody heavy chains from PCs isolated from HCs (purple), non-inflamed tissue from UC patients in remission (turquoise), less-inflamed tissue from UC patients with inflammation (green) and inflamed tissue (red). Sequences were stratified based on antibody isotype. Brackets indicate statistical significance using a two-sided t-test with \*\**P* ≤ 0.01, \*\*\**P* ≤ 0.001, \*\*\*\**P* ≤ 0.0001 and ns=non significant. **h,** Violin plots showing selection pressure (nucleotide replacement mutation/nucleotide replacement+silent mutation) in antibody heavy chain V-genes (top row), kappa light chain V-genes (middle row) and lambda light chain V-genes (bottom row). Data is stratified based on the disease state of the source tissue and antibody isotype as indicated. Brackets indicate statistical significance using a one-sided t-test with \**P* ≤ 0.05, \*\**P* ≤ 0.01, \*\*\*\**P* ≤ 0.0001 and ns=non significant.

**Extended Data Figure 4:**
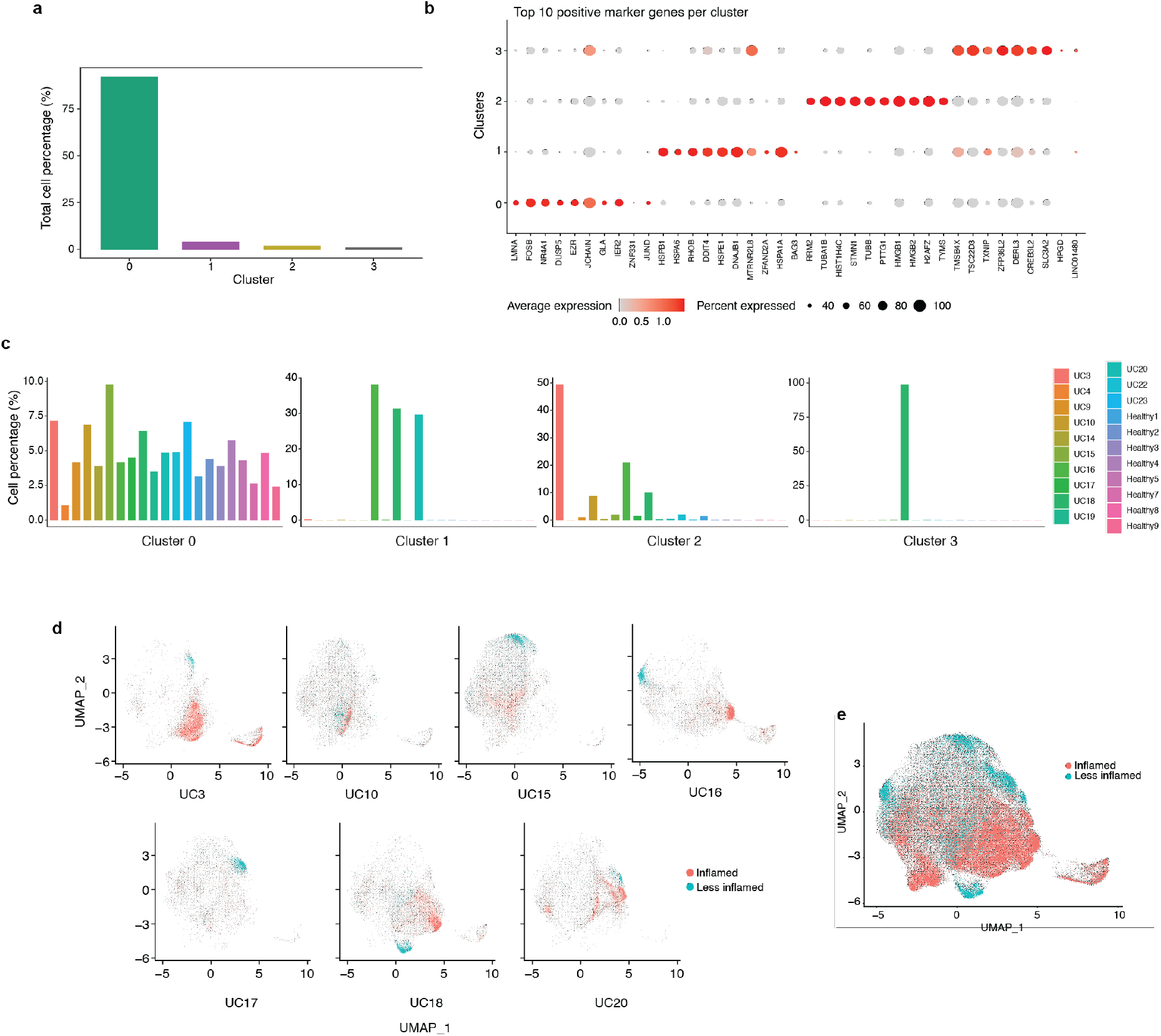
Transcriptional clusters of colon PCs. **a,** bar diagram displaying the fraction of all cells (in %) represented in each of the 4 transcriptional clusters. **b,** dot plot showing the relative expression of the top marker genes of each cluster (x-axis) across all clusters (y-axis). Each dot encodes both detection rate and average gene expression in detected cells for a gene in a cluster. Darker colour indicates higher average gene expression from the cells in which the gene was detected, and larger dot diameter indicates that the gene was detected in a greater proportion of cells from the cluster. **c,** bar diagram showing the representation of each of the 4 transcriptional clusters among all study subjects (x-axis) as indicated. Each plot represents the indicated cluster and the fractions add up to 100% in each plot. **d,** UMAP plots showing the cell embeddings colored by the inflammation status of the tissue the cells were isolated from for all subjects that had both inflamed and less inflamed tissue. **e,** UMAP plot showing all the cell embeddings shown in (**d**) but merged into one plot.

**Extended Data Figure 5:**
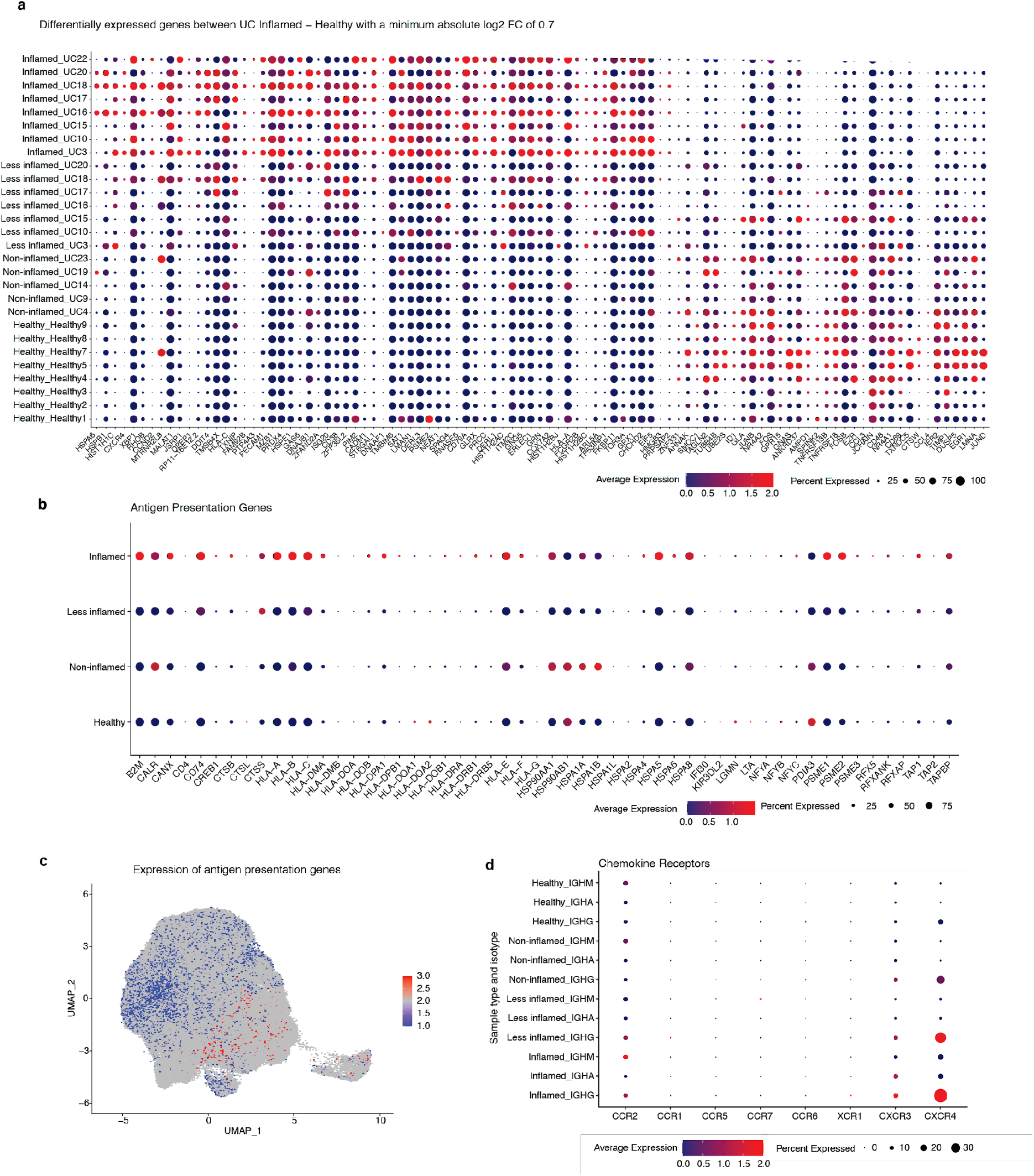
Genes that are higher expressed among PCs from inflamed tissue. **a,** Dot plot showing the relative expression of the significantly differentially expressed genes when comparing PCs derived from inflamed tissue with PCs from HCs (x-axis) across all subjects and disease states (y-axis) (pseudobulk DE analysis, FDR < 0.05). Each dot encodes both detection rate and average gene expression in detected cells for a gene in a cluster. As indicated, dark red color indicates higher average gene expression from the cells in which the gene was detected, and larger dot diameter indicates that the gene was detected in a greater proportion of cells from the cluster. **b,** Dot plot showing the relative expression of the antigen presentation genes (x-axis) in PCs derived from tissue in different disease states as indicated (y-axis). Color and size coding as in (**a**). **c,** UMAP plot showing the cell embeddings based on the transcriptome. Cells are colored based on their antigen presentation genes’ expression scores (**Methods**) where PCs highlighted in dark red show highest levels of expression. **d,** Dot plot showing the relative expression of chemokine receptor genes (x-axis) in PCs derived from tissue in different isotypes and disease states as indicated (y-axis). Color and size coding as in (**a**) and (**b**).

**Extended Data Figure 6:**
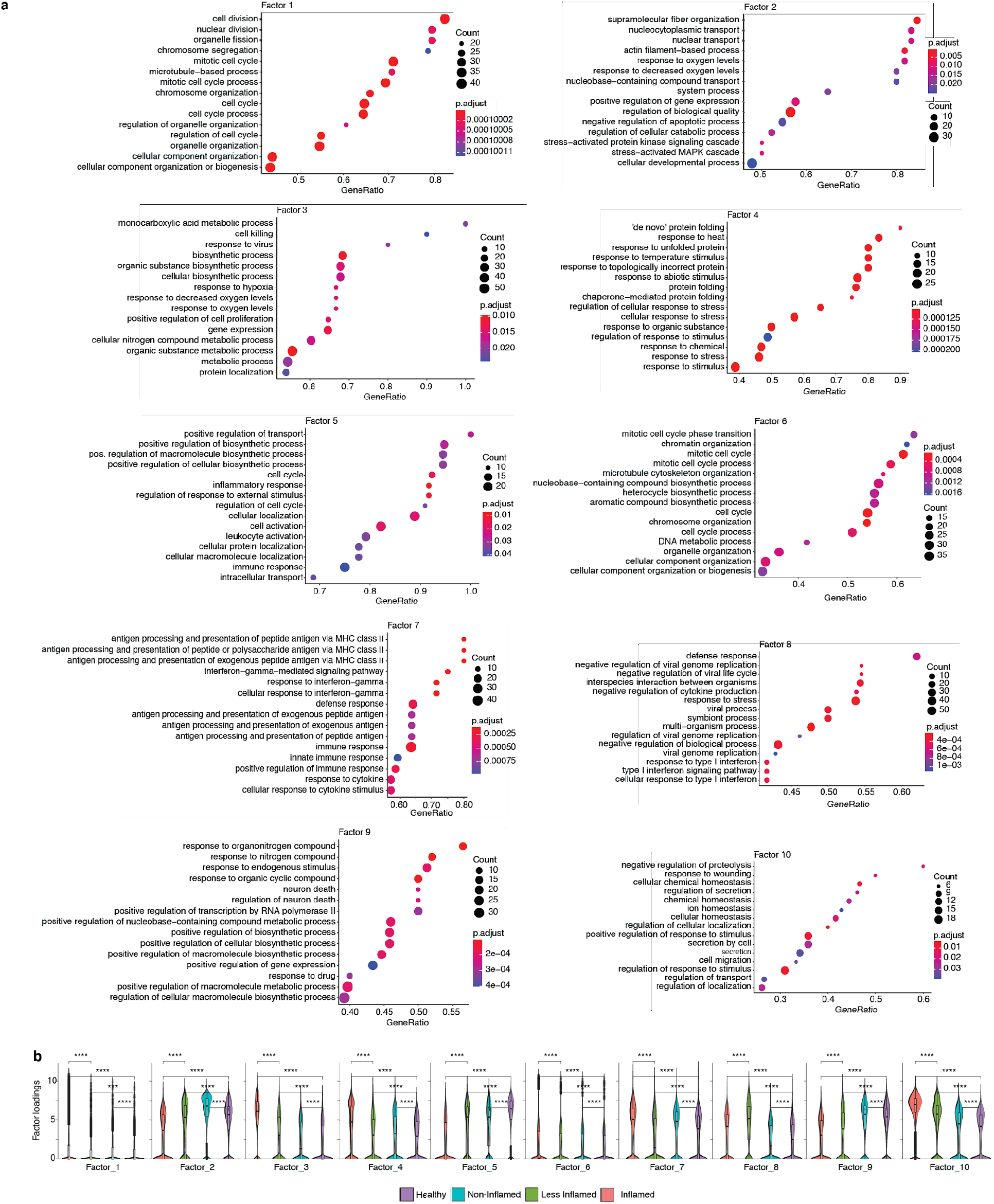
Latent factor analysis. **a,** Dot plots showing the top 15 gene ontology (GO) biological processes with the highest gene ratios in each of the 10 latent factors that were identified (**Methods**). The size of the dots represent the number of genes in the significant gene list associated with the GO term and the color of the dots represent the *P*-adjusted values. **b**, violin plots comparing the factor loadings between PCs from inflamed, less inflamed, non-inflamed and healthy control samples for 10 latent factors. \*\*\**P* ≤ 0.001 and \*\*\*\**P* ≤ 0.0001 using a one-sided t-test.

**Extended Data Figure 7:**
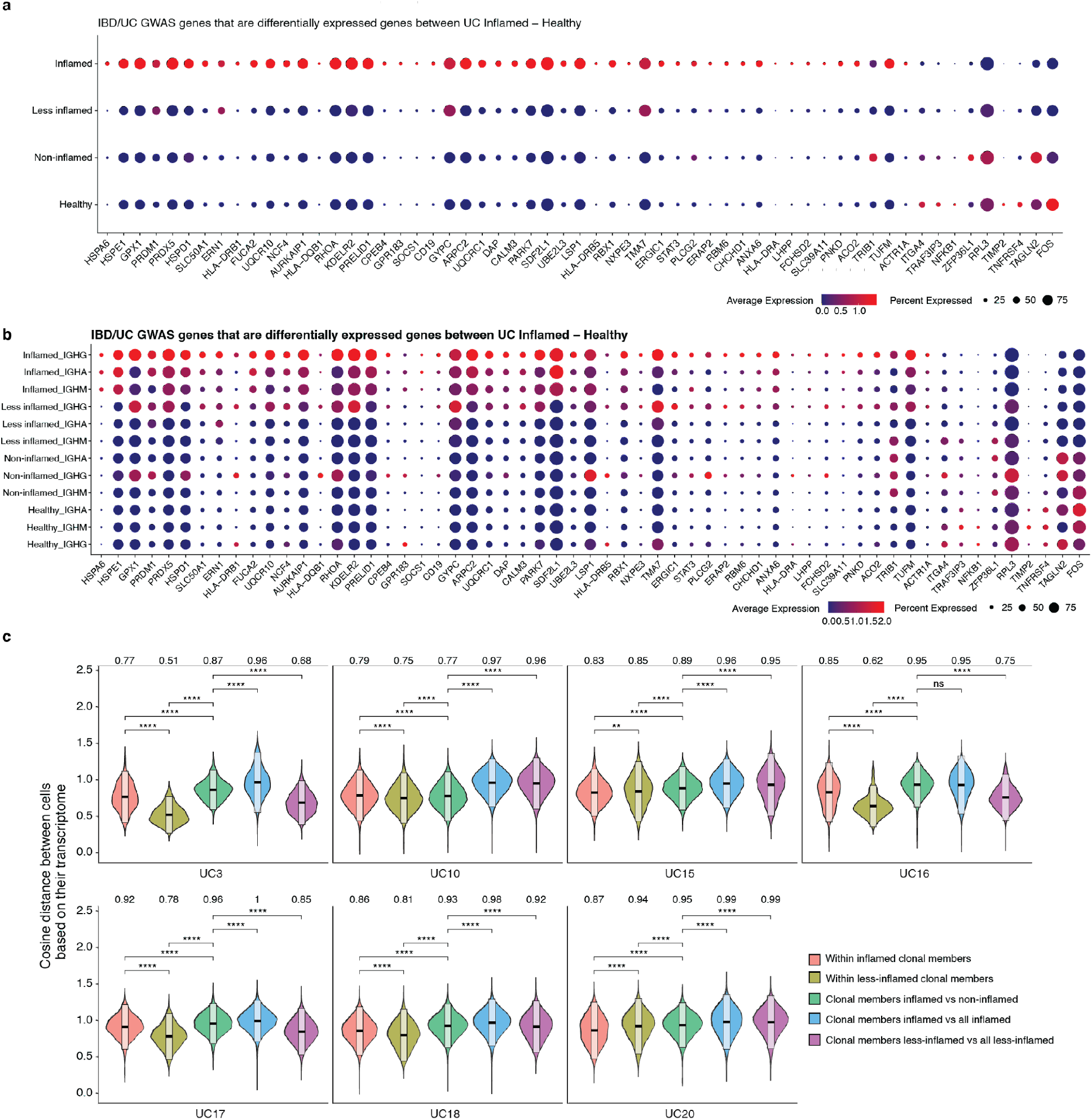
Expression levels of genes that are associated with IBD or UC through GWAS. **a, b,** dot plots showing the relative expression of the selected genes^23,24,25,26^ when comparing PCs derived from inflamed tissue with PCs from less inflamed, non-inflamed and HCs (y-axis). In (**b**) each disease state is stratified by antibody isotype. Each dot encodes both detection rate and average gene expression in detected cells for a gene in a cluster. As indicated, dark red color indicates higher average gene expression from the cells in which the gene was detected, and larger dot diameter indicates that the gene was detected in a greater proportion of cells from the cluster. (**c**) Comparison of the pairwise cosine distances between PCs based on their transcriptome for each subject with inflamed and less inflamed samples. Groups from left to right display the PC pair distance distributions between i) inflamed clone members, ii) less-inflamed clone members iii) inflamed and less-inflamed clone members iv) inflamed clone members and 100 randomly selected PCs of the donor that are inflamed and are not clone members v) clone members of less-inflamed samples and 100 randomly selected PCs of the donor that are less-inflamed and are not clone members.

**Extended Data Figure 8:**
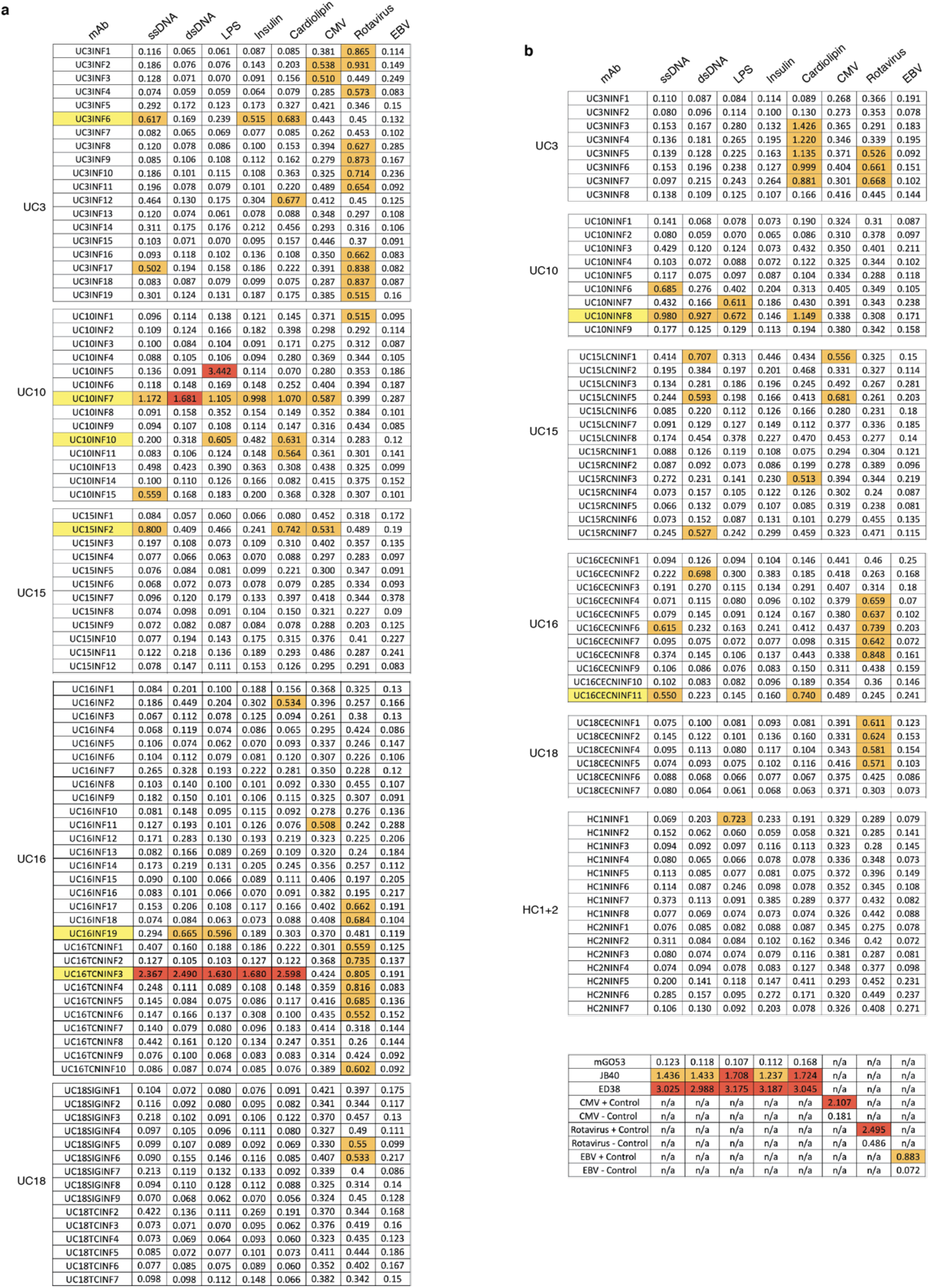
mAb reactivities in ELISA. Shown are the OD415 or OD450 values for each mAb at a concentration of 1ug/ml tested in ELISA for binding against single stranded DNA (ssDNA), double stranded DNA (dsDNA), lipopolysaccharide (LPS), cardiolipin, cytomegalovirus (CMV), rotavirus and Epstein-Barr Virus (EBV) (**Methods**). mAbs are sorted based on the donor and if they were isolated from inflamed (**a**) or non-inflamed (**b**) colon. Polyreactivity was defined as reactivity against at least 2 of the antigens ssDNA, dsDNA, LPS, insulin or cardiolipin as previously described^31, 34^. Polyreactive mAb names are colored in yellow. Orange color indicates low reactivity (OD=0.5-1.5) and red color indicates high reactivity (OD>1.5). Results for highly-, moderately- and non-polyreactive control mAbs ED38, JB40 and mGO53 are included as indicated^30^ as are positive and negative controls for CMV, rotavirus and EBV ELISAs (**Methods**).

**Extended Data Figure 9:**
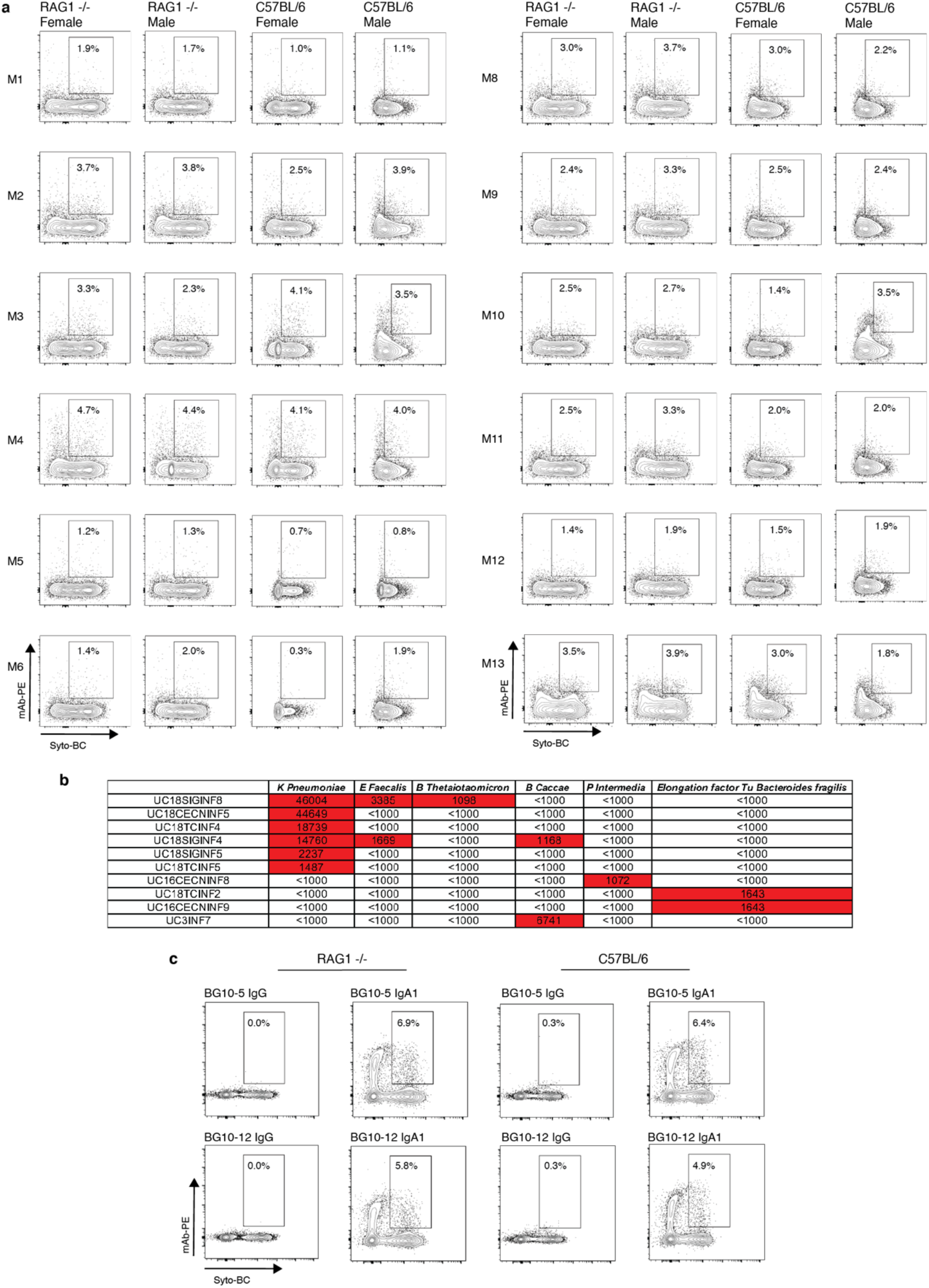
Binding of mAbs and mAb mixes to stool from RAG1-deficient and C57BL/6 mice and bacterial proteins and extracts. **a,** FACS plots display SYTO-BC (x-axis) and mAb mix staining (y-axis) of stool from female and male RAG1-deficient and C57BL/6 mice with the double positive population indicated through gating. Antibody binding was detected through a mouse anti-human IgG antibody coupled to PE (**Methods**). The composition of each antibody mix used is summarized in **Supplementary Table 5** and bacterial staining was conducted so that each mAb was present at a concentration of 10ug/ml. **b,** Table summarizing the fluorescence intensity values as measured on a GenePix 4000B imager (Axion) for the mAbs that showed binding in a screen of all 152 mAbs for binding to 50 bacterial lysates and antigens (**Supplementary Table 6** and **Methods**). Values above 1000 are considered positive and highlighted in red. **c,** FACS plots display SYTO-BC (x-axis) and mAb staining (y-axis) of stool from RAG1-deficient and C57BL/6 mice with the double positive population indicated through gating. SARS CoV-2 mAbs BG10-5 and BG10-12^43^ in IgG1 and IgA1 form were used for staining as indicated and their binding was detected with mouse anti-human IgG or mouse anti-human IgA antibody coupled to PE (**Methods**).

**Extended Data Table 1:**
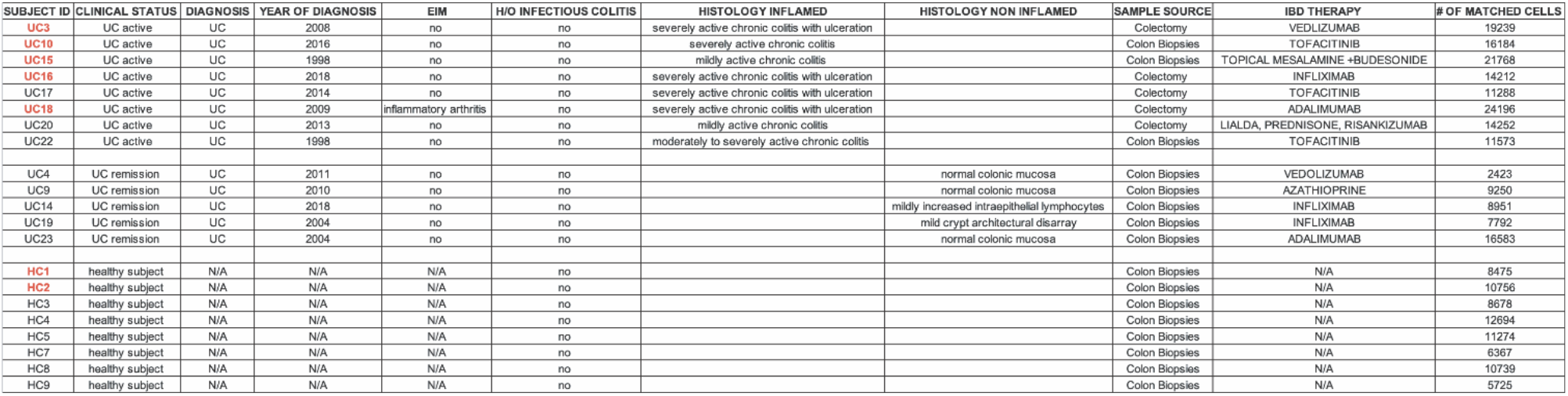
Patient characteristics. Shown are the subject IDs, clinical status, clinical diagnosis, year of diagnosis, presence of extraintestinal manifestations (EIM) of IBD such as inflammatory arthritis, dermatologic or ophthalmologic complications, history of infectious colitis, histology reports from the inflamed and non-inflamed colon areas sampled in our study (if available), the sample source and current IBD therapy as well as the number of matched VDJ and transcriptome cell profiles that were retrieved. F=female, M=male, UC=ulcerative colitis, HC=healthy control.

